# A comprehensive series of temporal transcription factors in the fly visual system

**DOI:** 10.1101/2021.06.13.448242

**Authors:** Nikolaos Konstantinides, Anthony M. Rossi, Aristides Escobar, Liébaut Dudragne, Yen-Chung Chen, Thinh Tran, Azalia Martinez Jaimes, Mehmet Neset Özel, Félix Simon, Zhiping Shao, Nadejda M. Tsankova, John F. Fullard, Uwe Walldorf, Panos Roussos, Claude Desplan

## Abstract

The brain consists of thousands of different neuronal types that are generated through multiple divisions of neuronal stem cells. These stem cells have the capacity to generate different neuronal types at different stages of their development. In *Drosophila*, this temporal patterning is driven by the successive expression of temporal transcription factors (tTFs). While a number of tTFs are known in different animals and across various parts of the nervous system, these have been mostly identified by informed guesses and antibody availability. We used single-cell mRNA sequencing to identify the complete series of tTFs that specify most *Drosophila* medulla neurons in the optic lobe. We tested the genetic interactions among these tTFs. While we verify the general principle that tTFs regulate the progression of the series by activating the next tTFs in the series and repressing the previous ones, we also identify more complex regulations. Two of the tTFs, Eyeless and Dichaete, act as hubs integrating the input of several upstream tTFs before allowing the series to progress and in turn regulating the expression of several downstream tTFs. Moreover, we show that tTFs not only specify neuronal identity by controlling the expression of cell type-specific genes. Finally, we describe the very first steps of neuronal differentiation and find that terminal differentiation genes, such as neurotransmitter-related genes, are present as transcripts, but not as proteins, in immature larval neurons days before they are being used in functioning neurons; we show that these mechanisms are conserved in humans. Our results offer a comprehensive description of a temporal series of tTFs in a neuronal system, offering mechanistic insights into the regulation of the progression of the series and the regulation of neuronal diversity. This represents a proof-of-principle for the use of single-cell mRNA sequencing for the comparison of temporal patterning across phyla that can lead to an understanding of how the human brain develops and how it has evolved.

---

The brain is the most complex organ of the animal body: the human brain consists of over 80 billion neurons^1^ that belong to thousands of different neuronal types and form ∼10^14^ synapses. Understanding the generation of this complexity in humans is an almost insurmountable problem. Researching the brains of simpler genetic model organisms as diverse as mice and flies provides windows into the underlying molecular mechanisms of how the production of neuronal diversity is achieved. This has highlighted the importance of two essential factors: the spatial location and age of neuronal progenitors.

Spatial information has long been acknowledged for its importance in patterning the dorsoventral and anteroposterior axes of animal bodies. Classic examples include patterning of the *Drosophila* embryo as well as the vertebrate spinal cord^2,3^. Morphogenetic gradients are converted into discrete spatial domains (e.g., the French Flag model^2^); in the nervous system, these domains express different transcription factors (spatial transcription factors - sTFs) and give rise to fate-restricted groups of neural stem cells, each of which can generate a unique subset of neuronal types^4,5^. There have been reports of orthologous transcription factors (TFs) acting as sTFs in mice and flies: for example, Drop (also called Msh – muscle segment homeobox)/Intermediate neuroblasts defective (Ind)/Ventral nervous system defective (Vnd) are expressed along the DV axis of the *Drosophila* neuroectoderm, and their mouse orthologs Msx1/Gsx1/Nkx2-2 are expressed in progenitor cells along the DV axis of the mouse neural tube at embryonic day 11.5^6^.

Temporal patterning describes the developmental trajectory neural stem cells follow that allows them to generate different neuronal types as they age^7,8^. It is a powerful mechanism that allows neural stem cells to produce different neuronal types in the correct order and stoichiometry. The first mechanism of temporal patterning in neuronal systems was described in the *Drosophila* ventral nerve cord (VNC) where a cascade of temporal transcription factors (tTFs) is expressed in embryonic neural stem cells (neuroblasts) as they divide and age^9–11^. It was later suggested that tTFs also contribute to the generation of neuronal diversity in different mammalian neuronal tissues, such as the retina^12–15^ and the cortex^16^. However, only a few tTFs have been discovered to play a role in both insects and vertebrates, such as Ikzf1 (a mouse ortholog of *Drosophila* Hunchback that is the first tTF in the fly VNC)^11,17^ that specifies young neural stem cells in the mouse cortex and retina^12,16^, as well as Pou2f1/Pou2f2^15^ and Casz1^13^ (the orthologs of the later VNC tTFs Pdm1/2 and Castor^11,17^) that specify older retinal progenitors.

The identification of tTFs in different neuronal systems has relied mainly on antibody availability (in *Drosophila)* or candidate genes from other systems (in mammals). This has hindered the identification of entire suites of tTFs and has made the evolutionary comparison of temporal series a piecemeal endeavor. The advent of single-cell mRNA sequencing (scRNASeq) allows for a comprehensive evaluation of the transcriptomes of neural stem cells of different ages and the exhaustive interrogation of all transcription factors for a potential role in temporal patterning. Recently, Telley et al^18^ profiled mouse cortical radial glia, as well as their immediate descendants, at different time points during development, offering an excellent resource to identify mouse tTFs. Although many genes were found to be dynamically expressed as the apical cortical progenitors age, a series of tTFs was not reported. Nonetheless, they discovered that neurogenesis of cortical excitatory neurons is governed by two orthogonal (*i.e.* independent) processes, specification and differentiation, where different neuronal identities are specified through time in radial glia while neuronal differentiation follows a precast and highly similar program.

The *Drosophila* optic lobe is an ideal system to address how neuronal diversity is generated and how neurons proceed to differentiate. It is an experimentally manageable, albeit complex structure, for which we have a very comprehensive catalogue of neuronal cell types. It consists of four neuropils, the lamina, medulla, lobula, and lobula plate. Meticulous work from the last decades have identified ∼100 cell types in the optic lobes based solely on morphological characters^19^. Recent work took advantage of elaborate molecular genetic tools, as well as scRNASeq, to expand the number of neuronal cell types to ∼200, based on both morphology and molecular identity^20–22^. Importantly, because the optic lobe processes visual information generated in each of the 800 unit eyes (ommatidia), it is formed of 800 similar circuits running in parallel, with many of the optic lobe cell types present in multiple copies in the brain (ranging from ∼5 to 800). This makes clustering of scRNASeq data and cluster annotation easier. Moreover, the neuroblasts that generate the medulla, which is the largest neuropil of the optic lobe, are formed by a wave of neurogenesis over a period of days, and they all progress through the same tTF temporal series^23^. This means that at any given developmental stage from mid third larval stage (L3) to the beginning of pupation, the neurogenic region contains neuroblasts at all stages of their development when each must express one of the tTFs (Figure 1A).

**Figure 1:**
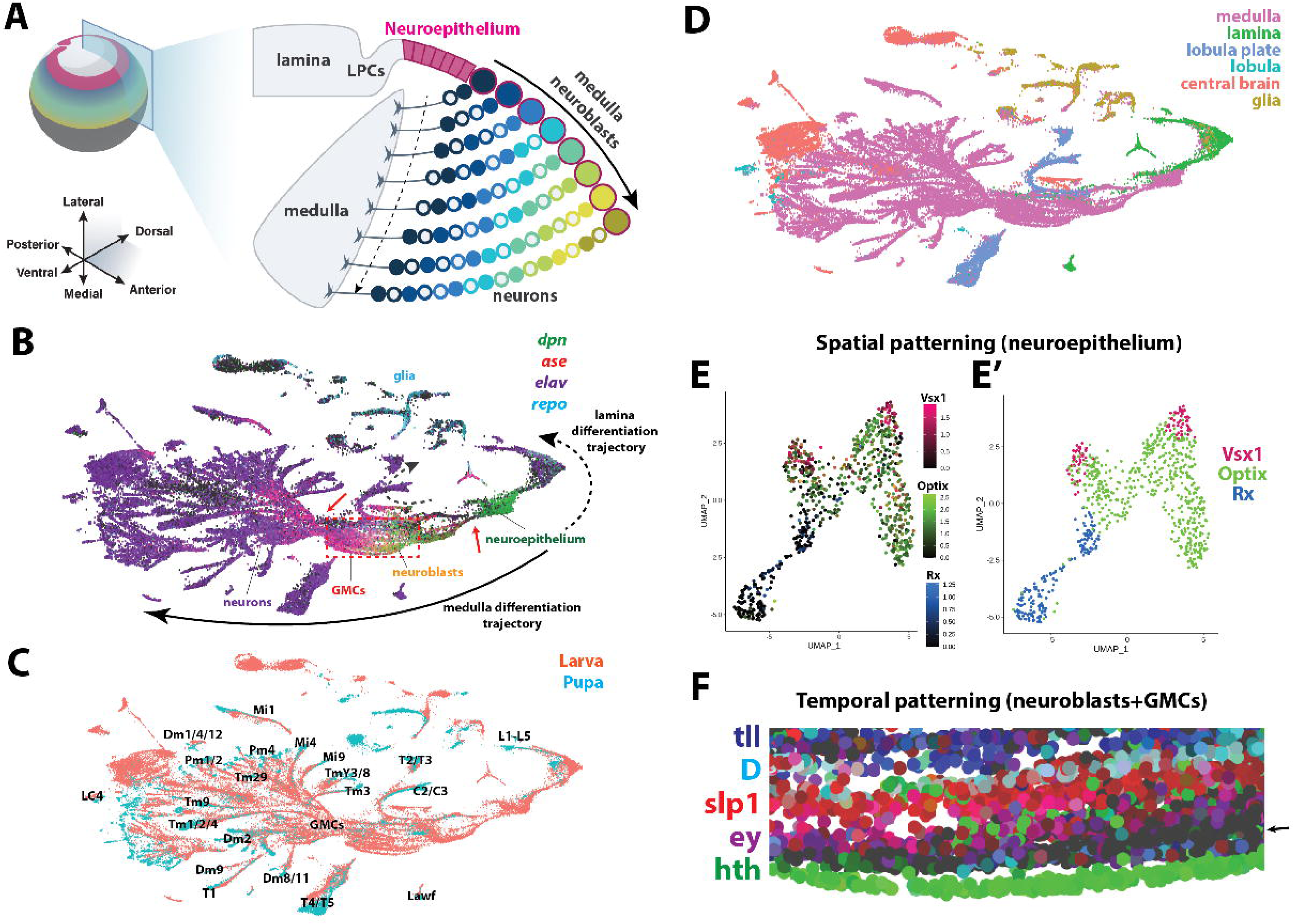
Single-cell sequencing of the *Drosophila* optic lobe. (A) Left: Schematic of the developing *Drosophila* optic lobe (colored) and central brain (dark grey) in the third larval stage. Right: Cross-section of the optic lobe. On the lateral side, the neuroepithelium (pink rectangles) is converted to lamina precursor cells (LPCs) that will form the lamina. On the medial side, neuroepithelial cells are gradually converted into neuroblasts (large circles) in a wave of neurogenesis. The neuroblasts, as they age, change their capacity to produce neuronal types (small circles). The solid arrow points from young, newly born neuroblasts to older ones. The dashed arrow points from young neurons to older ones. (B) UMAP plot of ∼50,000 single-cell transcriptomes from the developing optic lobes. The expression of *dpn, ase, elav,* and *repo* allows for the identification of the neuroepithelium (*dpn),* neuroblasts (*dpn* and *ase*), ganglion mother cells (GMCs – *ase*), neurons (*elav*), and glia (*repo*). The structure of the UMAP plot resembles the developing optic lobe with the lamina (dashed arrow) and medulla (black arrow) being generated from the two sides of the neuroepithelium. The IPC neurons (dashed arrow) are generated from different neuroblasts. The black arrows depict differentiation trajectories from neuroblasts to GMCs to neurons. The GMCs form an hourglass (left red arrow) with increased transcriptomic diversity when they are born and right before they divide into neurons. A similar hourglass is observed in the transition from neuroepithelium to neuroblasts (right red arrow). The dashed red box contains neuroblasts and GMCs that are highlighted in Figure 1F. (C) UMAP plot of single-cells coming from larval (pink) and pupal (cyan) developing optic lobes. The pupal cells fall on the tips of the neuronal branches that emanate from the center of the plot where the progenitors (neuroblasts and GMCs) reside. Notably, the pupal cells are annotated, which allowed us to identify both larval and pupal neuronal types. (D) Using this annotation, we were able to identify cell types that belong to the four optic lobe neuropils (lamina, medulla, lobula, and lobula plate), as well as central brain cells that were not removed during the dissections, and glial cells. (E) UMAP plot of the neuroepithelial cells. Left: Expression of the spatial transcription factors (Vsx1, Optix, and Rx) can be seen in largely non-overlapping clusters. Right: Semi-supervised clustering of the neuroepithelial cells and identification of the three spatial clusters (Vsx, Optix, and Rx). (F) Close-up (red dashed box of Figure 1B) of the UMAP plot showing neuroblasts and GMCs. The known temporal factors (*hth, ey, slp1, D,* and *tll*) are expressed in partially overlapping sets of neuroblasts (as has been shown experimentally) organized in the plot from top to bottom. *Hth* is expressed in the bottom row, followed by an empty row of neuroblasts that express a previously unknown temporal transcription factor (arrow), and then followed by *ey*, *slp1, D,* and *tll*-positive neuroblasts.

We performed scRNASeq of the developing *Drosophila* optic lobe at the time when neural stem cells (neuroblasts) divide to generate the ∼200 neuron types that compose this brain structure. We focused on the neural stem cell that form the main structure of the optic lobe, the medulla. This allowed us to identify most, if not all, tTFs that are expressed in medulla neural stem cells, as well as to describe the genetic interactions among them that allow the tTF series to progress. Although we can define a general rule that a given tTF activates the next tTF and represses the previous one, we uncovered an unexpected degree of complexity, in which different tTFs assume different roles in the timing of the progression of the series and the generation of neuronal diversity. We also showed that tTFs control neuronal identity by regulating the expression of downstream cell type-specific TFs. Finally, we describe the very first steps that a neuron takes to differentiate; we find that all neurons express terminal differentiation genes as early as L3 and they follow a similar route for differentiation, independent of their identity, and that this progression is conserved in the human brain.

## Single-cell transcriptomes recapitulate the structure of the developing optic lobe

The adult *Drosophila* optic lobes start developing at L3 from the lateral parts of the larval central nervous system. The medulla part of the optic lobe, which is by far the largest and most complex optic neuropil, is formed by medulla neuroblasts that are generated by a neurogenic epithelium called the outer proliferation center (OPC)^23^. Over a period of two days, the OPC is progressively converted into neuroblasts by a neurogenic wave that initiates medially and continues laterally until the entire epithelium is consumed. This process results in the generation of seemingly identical neuroblasts that produce neuronal types throughout optic lobe development, meaning that at any single point in time, there are medulla neuroblasts of different ages (young to old) (Figure 1A). This characteristic of medulla development provides a distinct opportunity to study neuroblast and neuronal trajectories in unparalleled detail since all developmental stages coexist in the same brain. To achieve that, we performed scRNASeq on optic lobes microdissected from the central brain using the Chromium system (10x Genomics). We obtained 49,893 single-cell transcriptomes from 40 L3 optic lobes. We used the Seurat v3 integration pipeline^24^ to remove batch effects between the ten different libraries that were generated (Extended Data Figure 1).

We used known markers to identify neuroepithelial cells (Shotgun, Shg^25^, and Deadpan, Dpn^26^), neuroblasts (Shg^25^, Dpn^27^ and Asense, Ase^28^), intermediate neuronal precursors, i.e. ganglion mother cells (GMCs – Ase^29^ and absence of Dpn), neurons (Embryonic lethal abnormal vision, Elav^30^), and glia (Reversed polarity, Repo^31^) in a UMAP^32^ plot (Figure 1B – Extended Data Figure 2A).

The OPC neuroepithelium generates two of the optic lobe structures: it is progressively converted from the medial side to give rise to medulla neuroblasts while its lateral side gives rise to the lamina precursor cells that will form the lamina^23^ (Figure 1A). Notably, medulla neuroepithelium, neuroblasts, GMCs and neurons were arranged in the UMAP following a progression that resembles the *in vivo* differentiation process (“medulla differentiation trajectory” (Figure 1B). Similarly, lamina neuroepithelium, lamina precursor cells, and lamina monopolar neurons were also arranged following a similar differentiation trajectory (“lamina differentiation trajectory”) but in the opposite orientation of that of the medulla, highlighting the similarities of the UMAP trajectories with the actual differentiation process in the brain (Figure 1B). Emanating from the GMCs, different neuronal branches emerge that appear to represent developmental trajectories of different neurons (Figure 1B). Lobula plate neurons are generated from the neuroblasts of the inner proliferation center (IPC) that are distinct from those of the OPC that generate the lamina and medulla. These neuroblasts and the neurons that are generated from the IPC follow a different trajectory in the UMAP plot (“IPC neuron differentiation trajectory”, Figure 1B) and will not be discussed further.

To verify whether these trajectories retain the information of time, as suggested by the progression of neuroblasts to GMCs to neurons, we merged the larval single-cell dataset with the early pupal stage 15 (P15) dataset that we had previously generated^21^. These P15 neurons fell at the tip of each of the neuronal branches, confirming that these branches indeed represent neuronal trajectories (Figure 1C). Importantly, since the P15 dataset is annotated, this allowed us to identify the neuronal types at the tip of the trajectories. We could in fact identify neurons from all the neuropils of the optic lobe (lamina, medulla, lobula, and lobula plate) in the larval UMAP, which together accounted for 85% of the dataset. The annotation of the different neuropils was confirmed by looking at known markers of the lamina (*dac, gcm, so, eya, sim*^33^) and lobula plate (*D, tll, acj6, dac*^34,35^) (Extended Data Figure 2B). The remaining cells included a small number of central brain neurons and neuroblasts that were retained when cutting off the optic lobe (Figure 1D).

We then looked at the expression of the known spatial and temporal TFs in the neuroepithelium and neuroblasts, respectively. The neuroepithelium is divided into three broad domains by the expression of three spatial factors (Vsx, Optix, and Rx)^5^. These spatial factors were expressed in largely non-overlapping subsets of the neuroepithelial cells (Figure 1E, Extended Data Figure 2C). We clustered the neuroepithelial cells and used Vsx1, Optix, and Rx expression in each cluster to assign them to a spatial domain (Figure 1E’). The number of neuroepithelial cells corresponding to the different domain correlated with their size: Optix represented the largest spatial domain (spanning ∼65% of the epithelium), followed by Rx (23%) and Vsx (12%). However, as previously shown by immunostainings, their expression was not maintained in neuroblasts and neurons (Extended Data Figure 2C).

The medulla neuroblasts express a series of five tTFs (Hth, Ey, Slp, D, and Tll) in a temporal manner^36^. tTFs showed a very distinct pattern in the UMAP plot: not only were they expressed in subsets of neuroblasts, but neuroblasts and GMCs were organized based on their age, with progenitors expressing a tTF positioned between those expressing the previous and the next one in the temporal cascade (Figure 1F): Hth was present in the bottom row of the cluster, followed by Ey, Slp, D, and Tll with partial overlap among them, similar to what is observed with immunostaining *in vivo*^36^. Interestingly, neuroblasts positioned between Hth and Ey were not expressing any of the known tTFs, as was expected from *in vivo* stainings^36^ (Figure 1F – arrow, discussed later)

In general, we observed that the UMAP plot recapitulated remarkably well what is happening in the developing tissue: there were two different axes of time in the UMAP, a vertical axis that represents neuroblasts progressing through their temporal series of tTFs and a horizontal axis that represents cell state and differentiation status (i.e., neuroblast to GMC to immature neuron to mature neurons). We observed two bottlenecks along the developmental axis (Figure 1B, red arrows), one when the neuroepithelium is converted into neuroblasts and one when the GMCs with different temporal identities converge transcriptionally before they diverge again towards separate neuronal trajectories. This might be due to the fact that gene transcription during epithelial to mesenchymal transition in the first case and before the terminal division of the GMCs in the second case are obscuring more specific identity features.

## A comprehensive temporal series of transcription factors in the developing medulla

Although they cover almost the entire life of neuroblasts, the existing tTFs were discovered from educated guesses and screening of available antibodies. There is also clear evidence that there are additional tTFs, as the existing TFs are not able to explain the entire neuronal diversity in the optic lobe and there are neuroblasts between the Hth and Ey temporal windows that do not express any of the five tTFs *in vivo*^36^. We could confirm that there are cells in the UMAP plot that express none of the known tTFs (Figure 1F - arrow). The larval scRNAseq dataset gave us the opportunity to look for all potential tTFs in an unbiased and comprehensive way. We isolated the cluster of medulla neuroblasts from the scRNASeq data and used Monocle^37^ to reconstruct their developmental trajectory. To confirm the accuracy of the trajectory, we looked at the expression of the known tTFs: Hth, Ey, Slp, D, and Tll. Indeed, as was already clear from the UMAP plot, these tTFs were expressed in the correct temporal order along the trajectory (Figure 2A – TFs in purple). We then examined the expression dynamics of all *Drosophila* TFs and identified 39 candidate tTFs that exhibited expression restricted to a temporal window. These fell into two distinct categories: 14 of them were expressed at relatively high levels and included the 6 known tTFs (Slp includes two genes, Slp1 and Slp2) (Extended Data Figure 3A), while 25 of them were expressed at lower and fluctuating levels along the trajectory (Extended Data Figure 3B). We tested the expression pattern of four of the 25 lowly expressed candidates (*ap, cut, gcm*, and *gem*) in the developing optic lobes using antibodies against Ap and the cut-Gal4, gcm-Gal4, and gem-Gal4 lines, but none were expressed in a temporal manner; therefore, we decided not to pursue these candidates further as their fluctuations likely represent noise. Moreover, although Klumpfuss (Klu) was previously suggested to be a tTF^38^, Klu mRNA was found to be continuously expressed throughout neuroblast life in our dataset. We then tested the protein expression of the eight newly discovered candidate tTFs in medulla neuroblasts (surface view - Figure 2B). Using antibodies against these potential tTFs and the already known ones, we verified that their protein expression was limited to a specific temporal window (Figure 2B’ and 2C). They are described below in their order of expression in neuroblasts.

**Figure 2:**
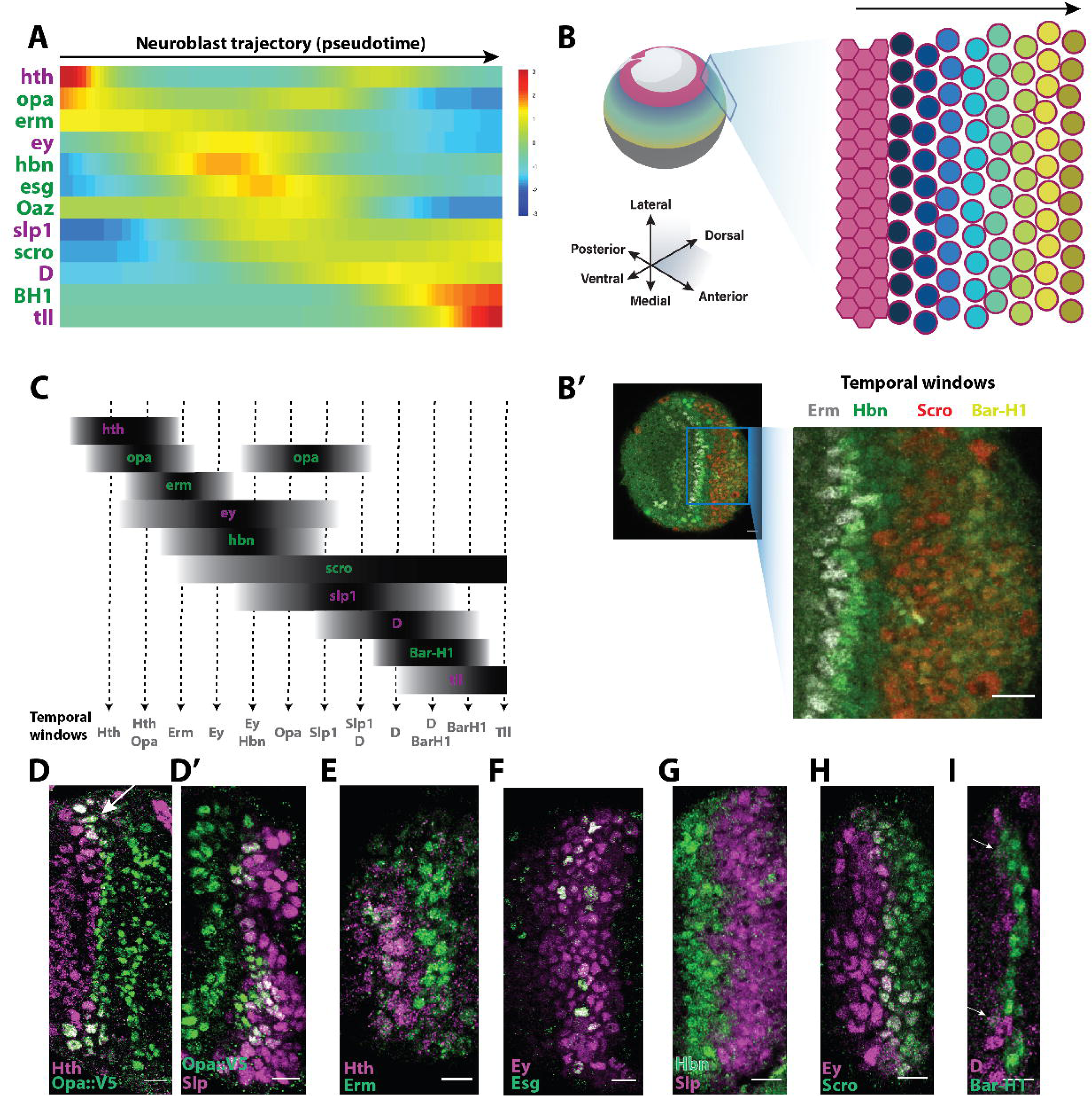
Newly identified temporal transcription factors are expressed temporally. (A) Temporal expression of known and candidate temporal transcription factors (y-axis) along the neuroblast developmental trajectory (x-axis, pseudotime), as generated using Monocle. Known temporal transcription factors are illustrated in purple, while newly identified candidate temporal transcription factors are shown in green. The colors represent scaled expression along the trajectory (red and yellow show expression, while cyan and blue absence of expression). (B) Left: Schematic of the developing *Drosophila* optic lobe (colored) and central brain (grey) in the third larval stage. Right: Surface (flattened) view of the neuroepithelium (pink) and neuroblasts. Young newly born neuroblasts are next to the neuroepithelium, while older ones are further away. (B’) Antibody staining (surface view) against four of the five new temporal transcription factors, Erm (white), Hbn (green), Scro (red), BarH1 (green). Left: The newly identified temporal transcription factors define new temporal windows, as illustrated in the entire developing optic lobe. Right: Close-up of the neuroblasts showing the four temporal windows: Erm (white), Hbn (green), Scro (red), and BarH1 (yellow). (C) Schematic of the expression of all tTFs in the ageing optic lobe neuroblasts. Dashed lines represent hypothetical divisions. With this suite of tTFs, a neuroblast can undergo ∼12 divisions with a distinct tTF identity (temporal windows). (D-I) Antibody stainings of newly identified temporal transcription factors (green) and previously known ones (purple) show that the candidate temporal transcription factors are indeed expressed temporally. (D-D’) Opa^62^ is expressed in two waves, one succeeding and partially overlapping (arrow) with the Hth window and one immediately before Slp. (E) Erm is expressed immediately after Hth. (F) Esg is expressed in a salt-and-pepper manner within the Ey temporal window. (G) Hbn is expressed before Slp1. (H) Scro is expressed immediately after Ey. (I) BarH1 is expressed after the D temporal window, slightly overlapping with it (arrows). Scale bar: 10um

**Opa** (Odd-paired) is expressed in two waves: it is first expressed in young neuroblasts immediately after and partially overlapping with the Hth temporal window (Figure 2D - arrow). Then its expression ceases before reappearing just before Slp (Figure 2D’ and Extended Data Figure 4A). **Erm** (Earmuff) immediately follows Hth (Figure 2E and Extended Data Figure 4B), partially overlaps with Opa and precedes (partially overlapping) Ey expression (Extended Data Figure 4C). **Esg** (Escargot) is expressed within the Ey temporal window, albeit in a salt-and-pepper manner, indicating that it is likely not a bona fide tTF as it is not expressed in all neuroblasts (Figure 2F and Extended Data Figure 4D). **Hbn** (Homeobrain) expression almost completely overlaps Ey in neuroblasts (Extended Data Figure 4E), right before Slp1 (Figure 2G and Extended Data Figure 4F). **Scro** (Scarecrow) expression starts immediately after Ey (Figure 2H and Extended Data Figure 4G), but it remains expressed until the end of the neuroblast divisions (Figure 2B’). **BarH1** is expressed after D (Figure 2I and Extended Data Figure 4H) and before Tll (Extended Data Figure 4I), partially overlapping both. Finally, in the absence of a functioning antibody, we tested the expression of **Oaz** using an Oaz-Gal4 line driving UAS-GFP. While based on the bioinformatic analysis it was expected to be expressed in young neuroblasts up to the Slp temporal window, it showed expression in all medulla NBs, potentially due to the perdurance of Gal4 in older neuroblasts (Extended Data Figure 4J).

Therefore, we could confirm that the predicted medulla tTF proteins (except potentially for Oaz) are expressed temporally in the developing optic lobe, defining new temporal windows as the neuroblasts progress through divisions (Figure 2C).

## Different tTFs assume different roles in the progression of the series

The known tTFs (except Hth) contribute to the progression of the series by activating the next tTF in the series and repressing the previous one^36^. While the TFs discovered above were expressed temporally, this does not imply that they actively participate in the progression of the temporal series. To test which of the newly identified tTFs were involved in the progression of the temporal series, we used mutant clones. We also drove UAS-RNAi constructs using the MZVUM-Gal4 line that is expressed in the Vsx1 domain^39^, in the central part of the medulla neuroepithelium (central Outer Proliferation Center) and its descendant neuroblasts and neurons. This allowed for the direct comparison with wildtype neuroblasts in neighboring control regions of the neurogenic region. Given the total number of tTFs in the medulla temporal series, we subdivided the temporal series into three broader units: early (between Hth and Ey), middle (between Ey and Slp), and late (between Slp and Tll) (Figure 3A - Extended Data Figure 5A).

**Figure 3.**
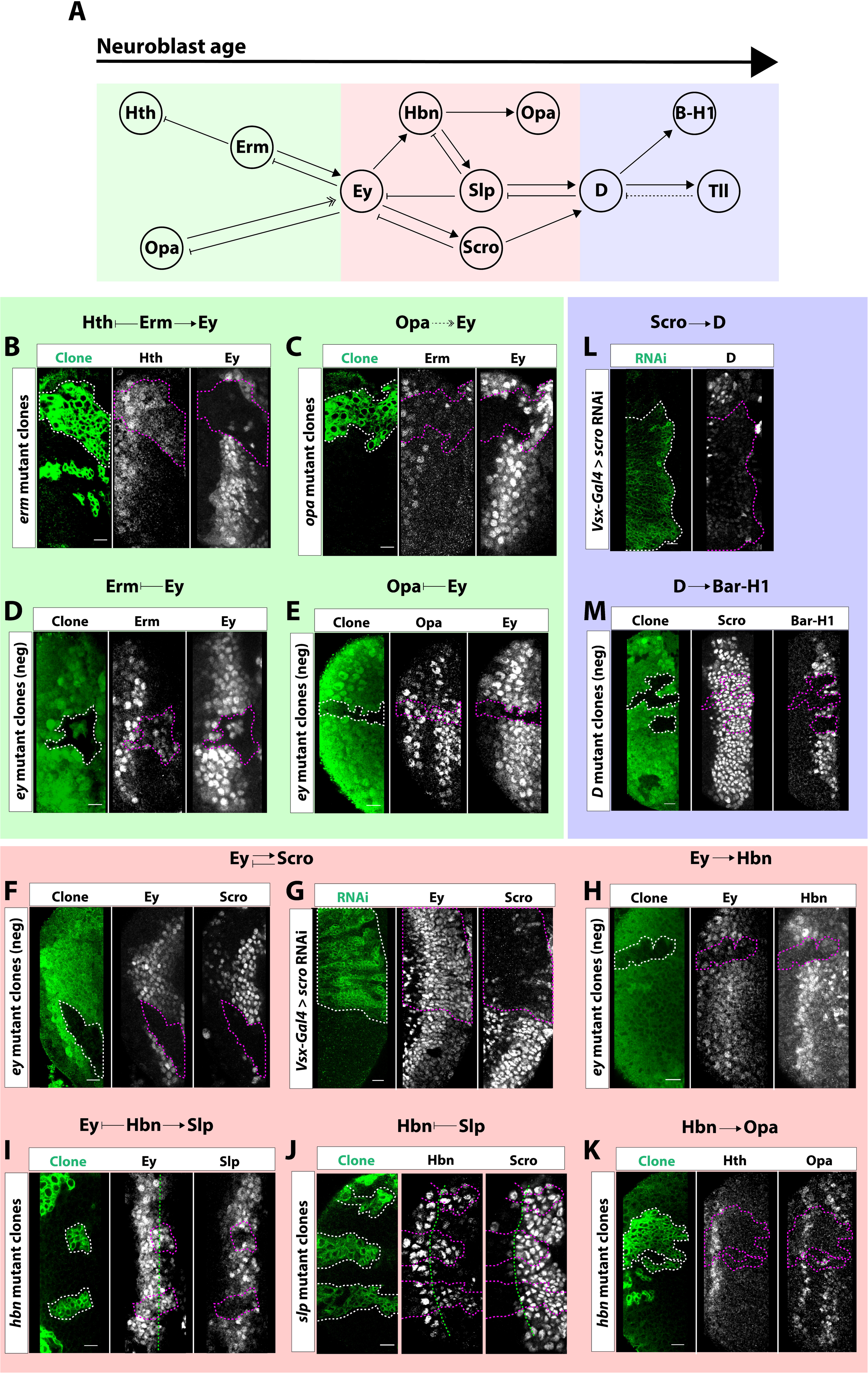
Complex genetic interactions between tTFs control the progression of the temporal series. (A) For clarity, the temporal series is subdivided into three units connected by two “hub” factors. The early unit (green box) is connected to the middle unit (red box) by Eyeless (Ey). The middle unit is connected to the late unit (blue box) by Dichaete (D). Within the early unit, we identified two new tTFs: Odd paired (Opa) and Earmuff (Erm). These factors help explain how neuroblasts progress from the Hth temporal window, which was not known. Within the middle unit, we identified three new temporal factors: Homeobrain (Hbn), Scarecrow (Scro) and Opa. Within the late unit we identified one new temporal factor: Bar-H1 (B-H1). (B) In *erm* mutant clones labeled by GFP (green), Hth is extended and Ey is lost. This indicates that Erm represses Hth and activates Ey. (C) In *opa* mutant clones (GFP: green), Erm is not affected and Ey expression is delayed, demonstrating that Opa only helps to time the expression of Ey. Sloppy paired (Slp) expression is also delayed in *opa* mutants (see Supp. Fig. 3). (D) Negatively labeled *ey* mutant clones (GFP-, Ey-) continue to express Erm, indicating that Ey represses Erm. (E) *ey* mutant clones (GFP-, Ey-) also continue to express Opa, showing that Ey also represses Opa. (F) The hub factor Ey both represses the entire network of early factors and also helps activate the second unit of temporal factors. In *ey* mutant clones (GFP-, Ey-), Scro expression is lost. (G) Conversely, Scro represses Ey, since cells expressing *scro* RNAi (GFP+: green; Scro-) continue to express Ey. Conversely, Slp and Hbn expression are not affected upon *scro* RNAi expression (see Supp. Fig. 3). (H) In addition to activating Scro, Ey activates Hbn, since Hbn expression is lost in negatively marked *ey* mutant clones (GFP-, Ey-). (I) *hbn* mutant clones (GFP:green) show that Hbn inhibits Ey and activates Slp. Taken together with our other results, this indicates that two parallel temporal series are activated by Ey (i.e., Ey/Hbn→Slp→D and Ey→Scro→D). (J) In *slp* mutant clones (GFP:green) Hbn expression is extended, showing that Slp inhibits Hbn. Conversely, Slp does not regulate the expression of Scro. (K) In addition to Hbn’s role in activating Slp to keep the temporal series progressing, Opa expression is lost in *hbn* mutant clones (GFP: green), indicating that Hbn activates the second Opa window. It remains unknown whether the second Opa temporal window helps regulate Hbn or Slp. (L) D acts as the second hub factor in the temporal series as both parallel temporal series converge to control D expression. In cells expressing *scro* RNAi, D expression is lost (see Supp. Fig. 3 for additional images). (M) In negatively marked *D* mutant clones (GFP-), Bar-H1 expression is lost but Scro expression is not affected, consistent with our observations that Scro is expressed until the very end of the temporal series (see Fig. 2). Previously published results showed that D activates Tll while Tll is sufficient but not necessary to inhibit D. Scale bar: 10um

### Early unit

Hth begins the temporal series: importantly, *hth* transcripts are present in the very first neuroblasts as well as in the neuroepithelium that has not yet been transformed into neuroblasts, indicating that its activation is likely regulated by upstream patterning events in the neuroepithelium. We had previously shown that Hth is not required for the progression of the temporal series as the next known tTF, Ey, is expressed normally in *hth* mutants^36^. Therefore, another overlapping factor must be responsible for activating Ey. In fact, we identified two factors that regulate the expression of Ey in different manners, Erm and Opa. Erm acts like the known tTFs as it is required to activate its next tTF, Ey and to inhibit the previous, Hth: In *erm* mutant clones, Ey is not expressed and Hth expression is expanded (Figure 3B). At the same time, Opa, which is co-expressed in the last Hth NBs, is required for the activation of Ey at the correct time: *opa* mutant neuroblasts have strongly delayed expression of Ey (Figure 3C), which leads to a delayed expression of the temporal series after Ey (Extended Data Figure 5E); Hth and Erm are unaffected in *opa* mutant neuroblasts (Extended Data Figure 5C). Once the Ey temporal window is initiated at the correct time by the combined action of Erm and Opa, Ey represses the expression of its activators, Opa and Erm: in *ey* mutant clones, both Erm (Figure 3D) and Opa (Figure 3E) are expanded to later temporal windows. Therefore, Erm is a tTF essential for the progression of the cascade while Opa contributes to the correct timing of expression of the next tTFs.

### Middle unit

We had shown that Ey activates and is then inhibited by Slp^36^. However, the developmental trajectory of neuroblasts uncovered a much more complex situation. First, we found that Ey also activates Scro and is inhibited by it: in *ey* mutant clones Scro expression was completely lost (Figure 3F), while when Scro was knocked down by RNAi, Ey remained expressed until the last division of the neuroblasts (Figure 3G). Moreover, Ey also activated Hbn and was inhibited by it: in *ey* mutant clones, Hbn expression was lost (Figure 3H), while in *hbn* mutant clones, Ey was extended to later temporal windows (Figure 3I). Then, Hbn allows the temporal series to progress by activating Slp as *hbn* mutant clones lacked Slp expression (Figure 3I). This suggests that the activation of Slp by Ey is mediated by Hbn. Slp then inhibits Ey^36^ and Hbn: *slp* mutant clones showed extension of Hbn into later temporal windows (Figure 3J). Finally, Hbn activates the second temporal window of Opa, as in *hbn* mutant clones, the second wave of Opa expression was absent (Figure 3K).

The complex genetic interactions that involve the activation of *ey* (temporal regulation by *opa* and regulation of expression by *erm*), as well as the fact that two genes (*Scro* and *slp*) are required for it to be repressed, indicate that Ey plays the role of a hub factor in the initiation and progression of the temporal series, as it integrates several signals before activating the expression of several downstream tTFs.

### Late unit

Finally, D requires both Slp and Scro to be expressed. We had previously shown that in *slp* mutant clones, D is not expressed^36^. Similarly, when Scro was knocked down by RNAi, D was not expressed in the neuroblasts (Figure 3L and Extended Data Figure 5G). Scro is therefore important for the progression of the series, as it is activated by Ey, which it then inhibits together with Slp; it then activates the expression of D and remains expressed until the end the neuroblast’s life. Once D is activated, it inhibits Slp and activates BarH1: in *D* mutant clones, BarH1 expression was lost (Figure 3M). At the same time, D activates Tll and Tll is sufficient to inhibit D, as was shown before^36^. However, Tll did not inhibit BarH1 (Extended Data Figure 5H).

We have thus been able to identify most, if not all, temporally expressed TFs in a developing neuronal system and to show that these tTFs participate in the progression of the temporal series. By exhaustively examining the genetic interactions between the new and old tTFs (Figure 3 and Extended Data Figure 5), we show that the temporal series is more complex than previously described and that not all tTFs play a similar role. While some tTFs directly inhibit the previous factor and/or activate the next (Erm, Hbn, Slp, and Tll), one does not participate in the progression of the temporal series (Hth) while others assume newly discovered roles: *(i)* During its first temporal window Opa regulates the timing of Ey expression (and, consequently, the timing of the rest of the temporal series), as elimination of Opa did not prevent the expression of Ey but delayed it significantly. *(ii)* Ey and D appear to be important hubs of the temporal series: Ey expression is activated by the combined action of Erm and Opa, and it requires two TFs to be repressed (Slp and Scro), indicating that it could be a checkpoint for the progression of the series, *i.e.* only when both Hbn and Scro are activated is the temporal series allowed to progress. Similarly, D needs both Scro and Slp to be expressed (Scro and Slp do not regulate each other – Extended Data Figure 5F), playing the role of a smaller hub, as well. *(iii)* Finally, Scro is the link between the two hub factors, being activated by Ey and activating D.

## Temporal transcription factors often remain expressed in neurons and regulate neuronal identity

Besides their participation in the progression of the temporal series, tTFs have been shown to also regulate neuronal identity either by being expressed in the neuronal subsets that are generated during their temporal window and acting as effector TFs (i.e. activating effector genes)^36^, or by activating the expression of downstream transcription factors^36,38^, which then regulate the expression of effector genes in the absence of the tTF.

We first looked at the expression of tTFs in the neuronal progeny in the scRNASeq data. Hth, Erm, Hbn, Ey, and D were expressed in largely non-overlapping neuronal clusters (Figure 4A-A’). Interestingly, although Hbn and Ey protein expression in neuroblasts coincides completely, the proteins are inherited by different neuronal types; for example, Hbn is expressed in Dm1, Dm3, Dm4, Dm9, and Dm12, while Ey is expressed in Mi4, Tm4, Tm29, and TmY5a. Moreover, Erm is expressed in neurons that also inherited Ey from the neuroblast, and D is co-expressed with Hbn in neurons that come from the Hbn temporal window (arrows); however, this represents a different function of these genes as their expression is not inherited from the neuroblasts. Finally, Scro was expressed mostly in late-born neurons (*i.e.* neurons that are born after the Ey temporal window), as expected by its expression in the neuroblasts (Figure 4A’’). On the other hand, Opa, BarH1, Slp, and Tll transcripts were not detected in any of the differentiating medulla neurons or glia. However, we could trace their expression in the GMCs at the root of different neuronal trajectories (Figure 4B), indicating that the GMCs that are born during these temporal windows do give rise to different neuronal types. Finally, the Tll window is known to give rise to glial cells and indeed the Tll^+^ GMCs were at the root of the glial trajectories (Figure 4B). To verify whether the neurons generated during the different temporal windows still expressed the tTF present in the neuroblast when they were generated, we immunostained larval optic lobes. If retained in neurons, the tTFs should form a laminar-like structure, in the order of appearance of the tTFs in the temporal series^40^. Indeed, all new temporal factors (Opa, Erm, Hbn, Scro, and BarH1) were expressed in neurons (Figure 4C). Interestingly, even the tTFs that ceased being expressed in the GMCs as mRNAs (i.e. *opa* and *BarH1*) persisted as proteins in the newly born neurons. This suggests that all tTFs, even those that are not actively expressed in neurons, play a role in controlling the expression of differentiation genes, or in relaying temporal identity information by activating downstream TFs.

**Figure 4:**
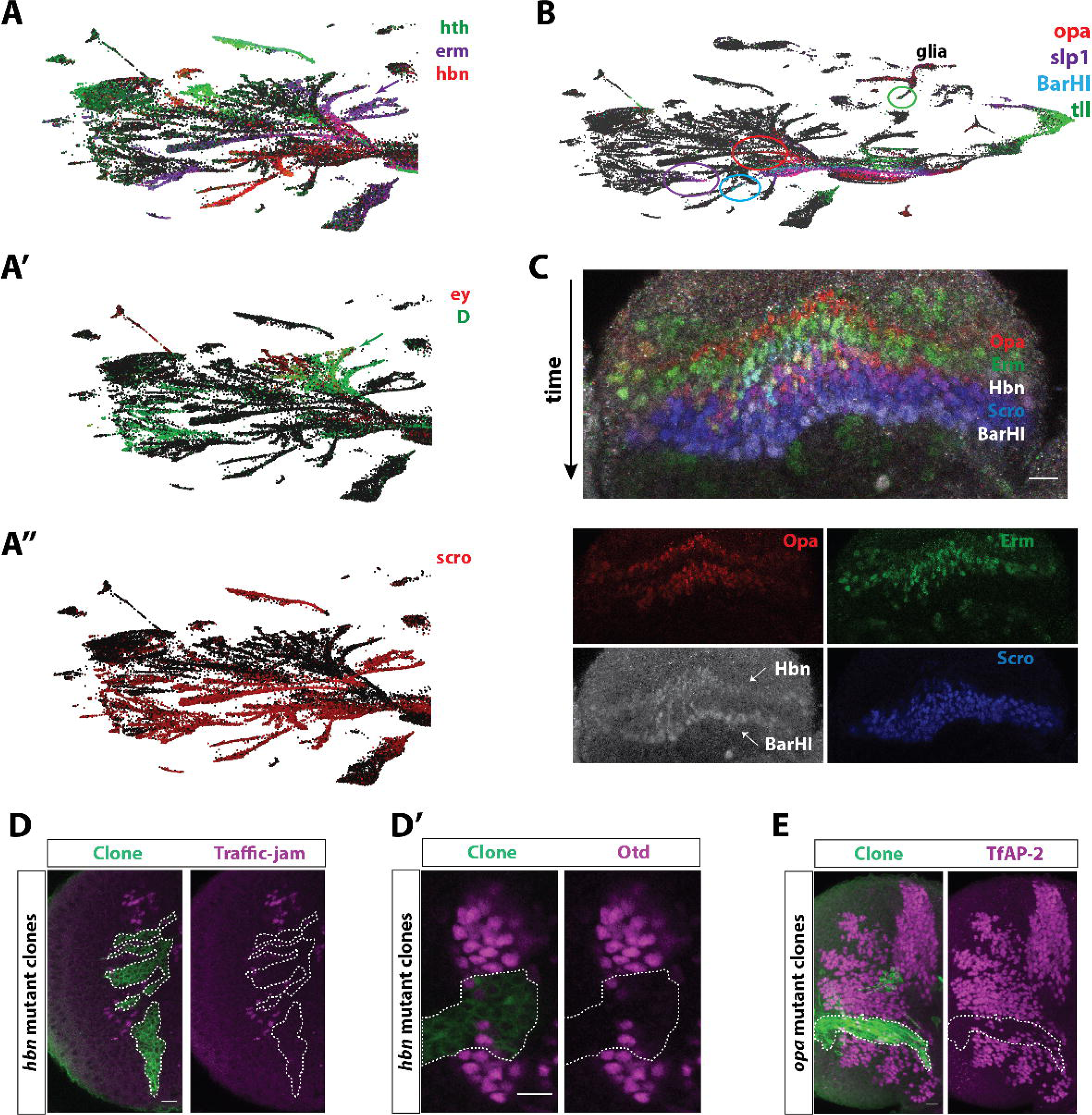
Temporal transcription factors are often maintained in neurons and regulate neuronal diversity. (A) UMAP plots showing the expression of all temporal transcription factors in the medulla neurons. (A) *Hth*, *erm*, and *hbn* are expressed in non-overlapping neuronal clusters. (A’) *Ey* and *D* are expressed in largely non-overlapping neuronal clusters. However, *D* is expressed in some of the progeny of the *ey* temporal window as has also been shown *in vivo*. (A’’) *Scro* is expressed in many neuronal types, as expected by its broad expression pattern in the neuroblasts. It is also expressed in neurons that express *hbn*, which suggests that *scro* is expressed in neuronal types that are generated earlier than the *scro* temporal window too. The arrows indicate the expression of *erm* and *D* in neurons coming from the Ey/Hbn temporal window. (B) UMAP plot showing the expression of *opa, slp1, BarH1*, and *tll*. *Opa*, *slp1*, *BarH1*, and *tll* transcripts are not detected in the neuronal progeny. However, they can be identified in GMCs in the root of neuronal branches (circles) suggesting that the GMCs that come from different temporal windows generate different neuronal types. (C) Expression of temporal transcription factors in neuronal progeny shows that the tTFs are expressed in the progeny of their respective neuroblast temporal window. Time is depicted by the arrow; neurons born from young neuroblasts are on the top of the Figure. Opa-positive neurons are born from young neuroblasts (red), followed by Erm neurons (green). Then, neurons expressing Hbn and Erm can be detected. (green-white). Finally, Scro-positive (blue) and Scro- and BarH1-positive neurons (blue-white) are born from older neuroblasts. Single-channel images can be seen in the bottom of the Figure. (D) Hbn is directly involved in the generation of neuronal diversity by regulating the expression of downstream transcription factors. *Hbn*-mutant MARCM clones (green) lack the expression of Traffic-jam (D) and Otd (D’) in the neurons. (E) Opa is also directly involved in the generation of neuronal diversity by regulating the expression of the downstream transcription factors TfAP-2 in some neurons. *Opa-*mutant MARCM clones (green) have fewer TfAP-2 positive cells (magenta) compared to the adjacent wild-type tissue. Scale bar: 10um

We then asked which neuronal types are generated from each temporal window; for this purpose, we used the expression of tTFs in neurons and GMCs described earlier to assign each neuronal type to a specific temporal window (Table S1). Except for the neurons described above that co-express Erm/Ey and D/Hbn, most tTFs are only expressed in neurons that come from their respective temporal window. We evaluated the assignments of the neurons to specific temporal windows by looking at neurons whose temporal window is already known, such as Mi1 (Hth temporal window), Tm3 (Ey temporal window), and Tm20 (Slp temporal window). We found that proximal medulla neurons Pm1 and Pm2 neurons come from the Hth/Opa temporal window, that Pm4 come from the Ey window, while all types of distal medulla (Dm) neurons come exclusively from the Ey/Hbn temporal windows. On the other hand, transmedullary neurons are generated throughout the neuroblast life (Opa, Ey/Hbn, Slp, and D temporal windows). We also looked for the expression of Hey, which is a Notch target^41^, to assess the Notch status of all neuronal types. We find 78 neuronal types to be Notch-OFF and 52 to be Notch-ON. This supports the idea that Notch-ON neurons ignore spatial cues and they are produced in all spatial domains of the OPC. As a consequence, Notch-ON neurons exhibit smaller diversity. This represents a unique resource to define the temporal origin and Notch identity of the cell types of the medulla part of the optic lobe and highlights the role of tTFs in regulating the generation of different neuronal types.

Finally, we asked whether knocking down the expression of the tTFs in neuroblasts could affect the expression of neuronal transcription factors. Bsh, Dfr, and Toy are transcription factors that are expressed in neurons that are born during a specific temporal window (Hth, Ey, and Slp temporal window, respectively). They are thus expressed in a laminar-like form and have been termed concentric genes^40^. The loss of *hth* in neuroblasts prevented the specification of Bsh-positive neurons, while in *ey* mutant clones, Dfr positive neurons were not specified, and in *slp* mutant clones, Toy neurons were absent^36,38^. We assessed the role of Hbn in neuronal specification by testing whether it is required for the expression of Traffic-jam and Otd that are expressed in neurons generated from the Hbn temporal window, as indicated by the trajectory of neurons in the UMAP plot. In *hbn* mutant clones, neurons expressing Traffic-jam and Otd (Figure 4D-D’) were no longer found. We also tested whether Opa was required for the generation of TfAP-2 positive neurons that are generated during both Opa temporal windows. In *opa* mutant clones, we found a significant reduction of TfAP-2 positive neurons compared to the adjacent wild-type tissue (Figure 4E). This shows that Hbn and Opa are regulating neuronal diversity not only by allowing the temporal series to progress (by activating the expression of Slp and timing the expression of Ey, respectively), but also by regulating neuronal transcription factor expression.

## The implementation of neuronal identity occurs very early in neuronal life and follows a fixed trajectory

*Drosophila* neurons are already specified at birth, based on the spatial and temporal identity of the neuroblasts from which they were born, as well as the Notch status of the neuron. However, it remains unclear how neurons proceed to differentiate. To study the very first steps of neuronal differentiation after specification, we initially focused on a well-studied and easily identifiable neuronal type, Mi1 (Mi1 is the only adult neuronal type in the medulla that expresses Bsh). We selected from the scRNASeq dataset of the L3 developing optic lobes the Bsh-positive clusters that correspond to Mi1 neurons at different levels of maturity, as well as the GMCs most closely linked to them in the UMAP plot (Extended Data Figure 6A-A’). We merged this dataset with the Mi1 clusters from pupal stages P15, P30, P40, P50, and P70, and used Monocle3^42^ to reconstruct their differentiation trajectory (Figure 5A). We used the expression of Ase in GMCs and Bsh in neurons to mark the beginning of the elongated trajectory at L3 and P15 (Extended Data Figure 6B-B’). At all later stages, the neurons formed a tight cluster with no clear trajectory. The larval cells were isolated from late L3 brains about 20 hours after neuroblasts started dividing and producing neurons. We therefore estimate that the L3 trajectory corresponds to the first ∼20 hours of neuronal life. We identified groups of genes (modules) that are co-regulated along the entire trajectory from L3 to P70 and searched for the GO terms enriched in each gene module (Figure 5B). The process of differentiation appears to follow a very specific path: Initially, at L3, cell cycle genes (False Discovery Rate - FDR=10^−20^) and DNA replication genes (FDR=10^−2^) were expressed, as expected from the division of GMCs. This was closely followed by an increase in genes involved in translation (FDR = 10^−141^), which may correspond to the increase in size of the neuron and the growth of neurites: GMCs that generate the neurons are the product of a heavily asymmetric neuronal division, which results in a large neuroblast and a small GMC that produces neurons that must grow in size. Then, genes related to dendrite development (FDR=10^−3^) and axon-guidance (FDR=10^−20^) started to be upregulated at late L3 until P15 when the neurons direct their neurites to the appropriate neuropils. These genes peaked around P15 to P30 before being downregulated at late pupal stages. Finally, genes important for neuronal function, such as neurotransmitter-related genes (FDR=10^−4^), cell adhesion molecules (FDR=10^−9^), synaptic transmission proteins (FDR=10^−4^), as well as channels involved in the regulation of membrane potential genes (FDR=10^−8^) started to be expressed as early as L3 before quickly reaching a plateau that was maintained until P15. Their expression then started to increase again until adulthood when the products of these genes support neuronal function (Figure 5B). This indicates that not only is neuronal identity specified during the first hours of neuronal development, but it is already implemented through genes involved in neuronal function very early (at least at the transcript level).

**Figure 5:**
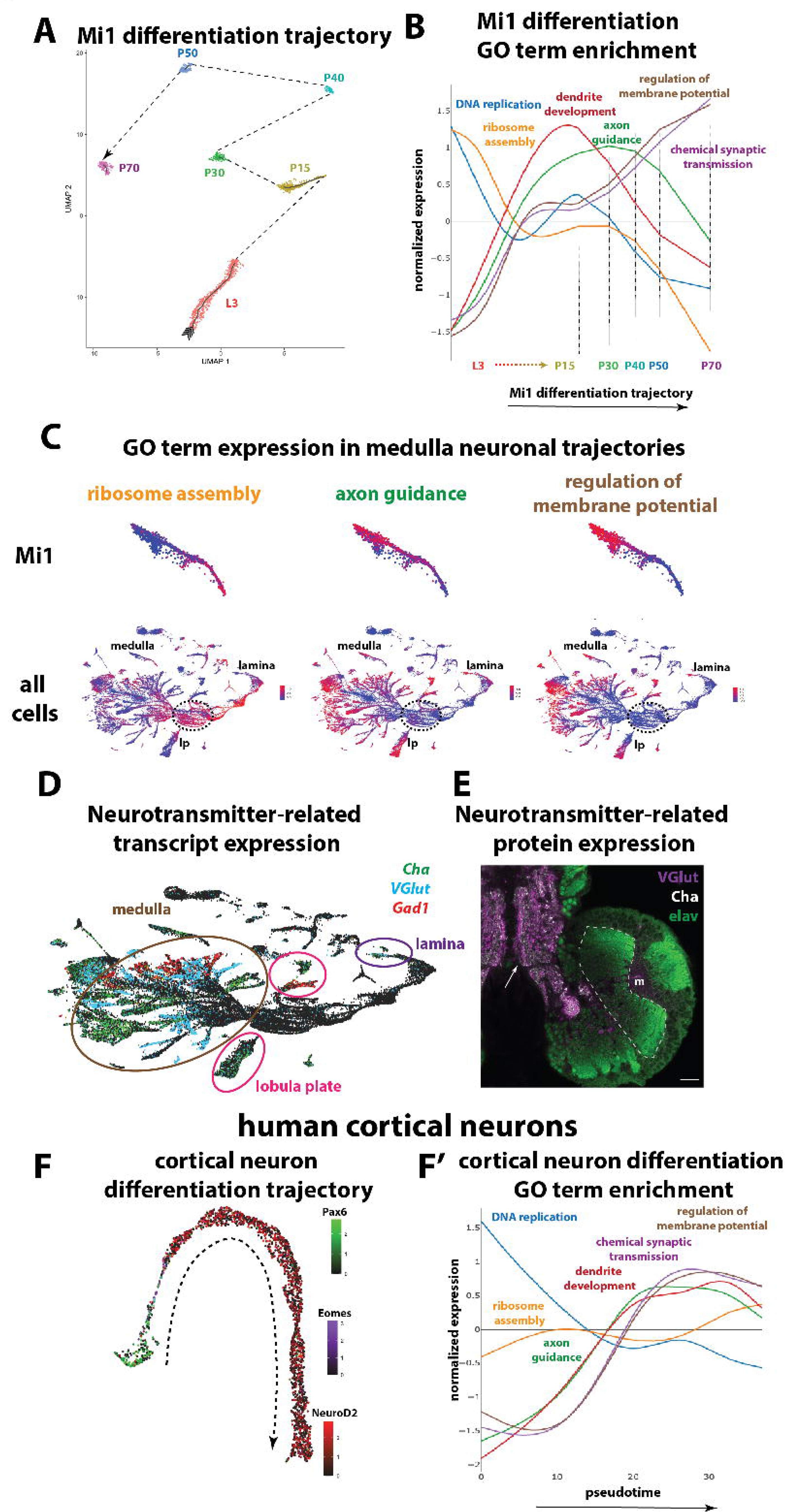
Neuronal differentiation in flies and humans. (A) UMAP plot of Mi1 cells at different stages of differentiation from L3 to P70. L3 and P15 trajectories are elongated depicting Mi1 cells of different ages. Transcriptomes are then synchronized (compact group of clusters) around P30. (B) Average expression of genes belonging to different Gene Ontology terms during the Mi1 differentiation trajectory. DNA replication is enriched in the beginning of the trajectory corresponding to the division of the GMCs that generate Mi1s. Ribosome assembly and translation-related terms are also expressed initially and persist into the newly born neurons to boost the translation machinery. Then, terms involved in neurite development and targeting, such as dendrite development and axon guidance, are enriched and peak around P15 and P30. Finally, neuronal function terms, such as regulation of membrane potential and neurotransmission start being upregulated as early as L3, reach a plateau around P15, before starting a drastic increase in P30 all the way to P70. (C) The differentiation route observed in Mi1s (top) is followed by all optic lobe neurons (bottom), including lamina and lobula plate neurons (lp). Average expression of genes belonging to three of the previous GO terms (ribosome assembly, axon guidance, and regulation of membrane potential) are depicted in the UMAP plots, which include cells from the L3 and P15 stages. Ribosome assembly terms are enriched (red) in dividing cells and young neurons. Genes involved in axon guidance start being expressed in neurons in the middle of their differentiation trajectory, while neuronal function terms, such as regulation of membrane potential, are expressed later towards the end of the trajectory, which represents the late L3 and P15 cells. (D) UMAP plot showing the expression of *Cha* (green), *VGlut* (blue), and *Gad1* (red), as markers for cholinergic, glutamatergic, and GABAergic cells, respectively, in medulla, lamina, and lobula plate. The transcripts can be detected in cells at the root of the trajectories, i.e. as soon as the neuron is specified. (E) Antibody staining of VGlut (purple), and Cha (white), in the developing *Drosophila* central nervous system at the third larval stage. Neurons can be observed using an antibody against Elav (green) in the optic lobe (dashed line), but there is no protein detected (although the transcript is present) in the neurons or the medulla neuropil (m). Conversely, expression of both VGlut and Cha can be detected in larval neurons of the ventral nerve cord (arrow). Scale bar: 20um (F) UMAP plot of 3,363 single-cell transcriptomes of the developing human cortex (gestational week 19). The trajectory from progenitors to neurons can be observed by the expression of Pax6 (apical progenitors), Eomes (intermediate progenitors), and Neurod2 (neurons). The dashed arrow depicts the differentiation trajectory. (F’) Average expression of genes belonging to the Gene Ontology terms that were described earlier over pseudotime. DNA replication is also enriched in the beginning of the trajectory when radial glia and intermediate progenitors divide. Ribosome assembly terms are not enriched at any point, contrary to the *Drosophila* neurons. Neurite development and targeting terms, such as dendrite development and axon guidance, are enriched in the beginning of the neuronal trajectory, followed closely by genes in involved in neuronal function, such as regulation of membrane potential and neurotransmission.

To evaluate whether this applies to all neurons or if each neuron follows a distinct differentiation trajectory, we aggregated and plotted the genes that belonged to the GO terms that were differentially activated during the Mi1 trajectory on the UMAP plot and observed that all optic lobe neurons followed the same differentiation trajectories, as indicated by the expression of the GO terms in all neuronal branches of the UMAP plot (Figure 5C). This differentiation trajectory was not only observed in the medulla, but also in the lamina and lobula plate neurons (Figure 5C).

As the expression of genes that should only be required very late was unexpected, we asked whether the transcript expression observed as early as late L3 was also translated into protein expression. We focused on neurotransmitter-related genes. We first verified that the correct neurotransmitter identity was indeed established as early as L3. We looked for the expression of ChAT, VGlut, and Gad1 (markers for cholinergic, glutamatergic, and GABAergic cell types, respectively) in the scRNASeq data and observed that they were expressed in the medulla in non-overlapping neuronal sets (Figure 5D). We confirmed using the adult scRNASeq data that we published recently^21^ that the neurotransmitter identity of each cell type at L3 was the same as in the adult (Table S1). We then stained late L3 brains with antibodies for choline acetyltransferase (ChAT) and the vesicular glutamate transporter (VGlut) in late L3. While we observed their expression in the mature neurons of the larval ventral nerve cord, we saw no expression in the developing optic lobe (Figure 5E). This suggests that transcribing the loci might be a means towards commitment to a specific neurotransmitter identity but that other factors prevent translation of these mRNAs.

In conclusion, we show that the differentiation process of *Drosophila* optic lobe neurons is fixed and independent of neuronal identity: Acquisition and implementation of identity are two consecutive processes, where the temporal and spatial information inherited from the neuroblasts specify the genes that are expressed, while the differentiation trajectory decides the timing of their expression. This agrees with recent data from the mouse cortex, where specification and differentiation were proposed to be two independent processes that occur mainly in different cell types (stem cells vs neurons) and where differentiation follows a precast path in all neurons, independent of their identities^18^.

## Common differentiation trajectory in *Drosophila* optic lobe and human cortical neurons

Understanding how neuronal differentiation occurs in human cortical neurons *in vivo* is necessary for the development of accurate *in vitro* differentiation protocols that can be used for neuronal replacement therapies^43^. We therefore wondered whether the differentiation trajectory we described in *Drosophila* optic lobe neurons was also implemented during human neuronal differentiation. We generated single-nuclear RNA-sequencing data from the developing human fetal cortical plate at gestational week 19. We used Monocle 3 to reconstruct their developmental trajectory (Extended Data Figure 6C). We could see a trajectory from radial glia progenitors to intermediate progenitors and postmitotic neurons, as indicated by the expression of Pax6^44,45^ in radial glia, Eomes^45^ in intermediate progenitors, and Neurod2^46^ in neurons (Figure 5F). We then performed differential expression analysis and identified gene modules that were co-regulated along the trajectory. We performed Gene Ontology analysis and observed a remarkable similarity to *Drosophila*; the first two modules were enriched in genes involved in cell proliferation (FDR=10^−3^) and DNA replication (FDR=10^−44^) while the third module was enriched in axon guidance (FDR=10^−7^) and dendrite development (FDR=10^−6^), before the activation of genes involved in regulation of synapse organization (FDR=10^−11^) and cell adhesion (FDR=10^−8^). Finally, the last module showed enrichment in neuronal activity-related GO terms, such as calcium-dependent exocytosis (FDR=10^−2^) (Extended Data Figure 6C’). To directly compare the human and *Drosophila* neuronal differentiation trajectory, we plotted the expression of the GO terms that were expressed at different stages of the differentiation trajectory in *Drosophila* (Figure 5B) on the human cortical differentiation trajectory (Figure 5F’). We observed very similar dynamics with an initial expression of DNA replication genes in radial glia and intermediate progenitors, and then the upregulation of axon guidance genes in neurons, before the expression of functional genes involved in synapse formation and function, and in action potential. The main difference that we observed was the absence of enrichment for ribosome assembly and translation-related GO terms at early stages. This could potentially be explained by the slower development of human neurons that leads to a slower increase in size compared to *Drosophila*, as well as the fact that the division of the radial glia is more symmetric in terms of size^47^ than the asymmetric division of the optic lobe neuroblasts. Despite this difference, these results show that the neurons follow a similar differentiation trajectory in *Drosophila* and humans that can be either attributed to convergence or to their common origin.

## *Drosophila* tTF expression in mouse cortical radial glia

While similarities in spatial patterning between vertebrates and flies has long been noted^3,6^, it is not clear how temporal patterning has evolved^48,49^. Sporadic evidence suggests that temporal factors identified in the *Drosophila* ventral nerve cord neuroblasts pattern stem cells that generate the mouse retina and even the cortex^12–16^. To address the similarities and differences between *Drosophila* neuroblasts and mouse cortical radial glia, we probed for the expression of known *Drosophila* tTFs a recently published scRNASeq dataset from the mouse cortex, where radial glia at different stages of development (E12-E15) were sequenced^18^. We first looked for the orthologs of the known tTFs of the *Drosophila* optic lobes described earlier, as well as in this study. None of the medulla neuroblast tTFs were expressed in strict temporal windows in ageing radial glia between day 12, where they produce deep cortical layers, and day 15, when more superficial layers are generated (Extended Data Figure 7A), with the exception of the Ey ortholog, Pax6, which was enriched in older progenitors. Foxg1, an ortholog of Slp was previously described to be enriched in young radial glia ^49,50^; while it is slightly reduced in older radial glia in this dataset, this reduction was not statistically significant. We also looked at the *Drosophila* orthologs of the mouse TFs that were described to be expressed in a temporal manner in the mouse CNS^51^; none of them were found to be tTFs in our trajectory analysis. The lack of conservation of a common temporal series of transcription factors *sensu stricto* between flies and mice shows that the acquisition of the specific temporal series occurred later and independently in each phylum, which is consistent with the several cascades of tTFs observed in various brain structures, such as the ventral nerve cord^11^, Type II central brain neuroblasts^52^ or optic lobe neuroblasts.

## Discussion

We present here a comprehensive series of transcription factors that temporally pattern a developing neural structure. We took advantage of the unique structure of the developing *Drosophila* optic lobe and generated detailed trajectories of neural stem cells starting from the time they are born to the time they terminally differentiate. All known tTFs were confirmed using in this approach; moreover, all the candidate tTFs that were identified computationally were verified experimentally, thus providing a proof-of-principle for the combination of scRNASeq and trajectory inference that can be applicable to most other neuronal tissues in different animals that lack the genetic toolkit of *Drosophila*. We show that most tTFs are expressed in overlapping windows creating a combinatorial code that differentiates neural stem cells of different ages and therefore provide them with the ability to generate diverse neurons after every division. We conservatively assigned them into 12 distinct temporal windows (Figure 2C), which, when integrated with spatial patterning (5 spatial domains) and the Notch binary cell fate decision (2 outcomes – Notch^ON^ and Notch^OFF^), can explain the generation of ∼120 cell types (12 times 5 times 2), which is close to the entire neuronal type diversity of the *Drosophila* medulla.

We also identified the regulatory interactions between these tTFs. Importantly, we show that not all tTFs function in the same way for the progression of the temporal cascade: while several tTFs directly control the progression by activating the next tTF and repressing the previous one, others (Ey and D) play important hub roles in this progression, by integrating both activating and inhibitory signals from several tTFs. The complex nature of the temporal series is likely a product of a complex evolutionary process that led to it, which will be very interesting to untangle in the future by comparing to other arthropod optic lobe temporal series.

We further show that the tTFs are not only important for the progression of the series but are also active contributors to the production of different neuronal types. Most of the tTFs are expressed in neuronal progeny, which suggests a role in establishing neuronal identity. They also regulate the expression of neuronal transcription factors, such as Bsh, Dfr, Toy, Traffic-Jam, Otd, and TfAP-2. These transcription factors are expressed in neurons that are born during a specific temporal window and represent the “business end” of temporal patterning, i.e. the TFs downstream of the tTFs that regulate terminal neuronal features. Importantly, these experiments allowedus to associate different neuronal populations with their temporal window of origin: by analyzing the expression of the tTFs expressed in these clusters and some of their downstream TFs, we were able to assign each of the different clusters of the optic lobe to a given temporal window, which is a unique resource to study these neurons (Table S1).

We also provide a detailed transcriptomic description of the first steps in the differentiation trajectory of a neuron. We find that all optic lobe neurons follow a similar differentiation program once they become postmitotic consisting of four main steps: *(i)* neuronal growth *(ii)* axon guidance, *(iii)* commitment to neurotransmitter identity and expression of cell adhesion molecules, and *(iv)* expression of genes encoding proteins that participate in mature neuronal function. Interestingly, all these steps are initiated within the first 20 hours of neuronal life, approximately 2-4 days before most of these proteins can fulfil their function. Why these genes are expressed so early remains unknown, but we hypothesize that it may be a way for neurons to commit to a specific identity. This path is taken by all neuronal types, independently of identity and of the actual genes that are expressed (*e.g.* neurotransmitter identity is established at the same relative time, independent of whether the neuron is cholinergic or glutamatergic) which is reminiscent of neurogenesis in vertebrates, as we show by analyzing single-nucleus sequencing data from the developing human cortex. This shows that, while specification mechanisms decide which genes are going to be expressed, the timing of the expression of genes of a specific function is preset, and this attribute of neurons is conserved between mammals and flies. Importantly, this highlights that understanding the mechanisms of neuronal differentiation in flies can generate insight for the equivalent process in humans, which is necessary for the generation of *in vitro* neuronal differentiation protocols^53^.

Finally, we probed a scRNASeq dataset of mouse radial glia for the expression of the optic lobe tTFs. None of the *Drosophila* tTFs are expressed in strict temporal windows; only Pax6 was found to be expressed in a gradient being enriched in older radial glia, while Foxg1 was slightly elevated in younger ones. Notably, the regulatory interaction between Ey/Pax6 and Slp/Foxg1 is potentially conserved between flies and mice^54^. Interestingly, the RNA binding protein Imp/Igf2bp1 is expressed in a gradient, being enriched in young progenitors in both *Drosophila* central brain neuroblasts and in the mouse cortical radial glia^55,56^ (Extended Data Figure 7B). Moreover, Bach2 (which has been suggested to be an ortholog of Chinmo^57^) is expressed in very young radial glia, but more importantly in neurons that come from the high-Igf2bp1 radial glia (Extended Data Figure 7B), This is reminiscent of the expression of its reported ortholog, Chinmo, that is also expressed in young neurons of the fly central brain and mushroom body under the regulation of Imp. This suggests that a similar temporal program may exist between radial glia and *Drosophila* neuroblasts that regulates neuronal identity.

The absence of strict temporal windows in mouse cortical apical progenitors was also illustrated by a recent paper that identified shared tTFs in the mouse CNS. These TFs were only divided in two broad categories, early *versus* late^51^. This supports the idea that tTF series with defined temporal windows, where tTFs count time and switch fate in response to a purely transcriptional network, can only operate in short-lived stem cells, like those in the *Drosophila* optic lobes that live for approximately 20 hours. On the other hand, gradients could be more adequate to count time in long-lived stem cells, such as the mouse radial glia that live for more than 5 days^58^.

Recent studies have highlighted key aspects of the evolution of the “nuts and bolts” of a functional neuron, such as the molecules of the pre- and post-synaptic machinery^59^ (i.e. how neurons evolved). On the other hand, other studies have focused on the evolutionary history of whole brain regions, such as the claustrum^60^, the hippocampus, the amygdala^61^ etc. The comprehensive comparison of the neuronal specification mechanisms, such as temporal patterning, will offer insight into the evolution of development of specific neuronal types. Only by combining all three different levels, evolution of neurons, neuronal types, and neuronal structures, will we be able to understand how the most complex organ of the human body evolved.

## Methods

### Genetics

To generate MARCM clones, crosses were kept at 25 °C and were heat-shocked for one hour at 37 °C four days before dissecting wandering L3 larvae. For RNAi experiments, MZVUM-Gal4 (Vsx-Gal4) flies were crossed to flies carrying the RNAi construct; the crosses were kept at 25 °C before dissecting wandering L3 larvae. The crosses are indicated below:

<u>*hth* RNAi:</u> *MZVUM-Gal4; UAS-CD8.GFP;* flies were crossed with *;hth-RNAi;* flies
<u>*scro* RNAi:</u> *MZVUM-Gal4; UAS-CD8.GFP;* flies were crossed with yv;; *scro-RNAi* flies
<u>*erm*- MARCM clones:</u>; erm1, FRT40A/CyO,act-GFP; flies were crossed with UAS-CD8GFP, hs-flp; FRT40A, tub-Gal80; tub-Gal4/TM6B flies.
<u>*opa*- MARCM clones:</u> *;;FRT82B - opa(null)/TM6B* flies were crossed with *yw, hs-flp, UAS-GFP;; tub-Gal4, FRT82B, tub-Gal80/TM6C* flies.
<u>*ey-* MARCM clones:</u> *yw,hsflp122; +/(Cyo); FRT80B/TM6B; ey[j5.71]/In(4)* flies were crossed with *yw, hsflp122; +/cyo; FRT80B ey-rescue (y+) ubiGFP/TM6B; ey [J5.71]/In(4)* flies.
<u>*D*- MARCM clones:</u> *yw; If/Cyo; D[87],FRT2A/TM6B* flies were crossed with *yw, hsflp; if/cyo; FRT2A, ubi-nlsGFP/TM6B* flies.
<u>*hbn* MARCM clones:</u> *FRT42B(G13), hbn15227* flies were crossed with *yw, hs-Flp; FRT42B(G13), tub-Gal80/CyO, act-GFP; tub-Gal4, UAS-CD8GFP/TM6,Tb,Hu* flies.
<u>*slp*- MARCM clones:</u> *yw, hsflp122; slp[s37a],FRT40A/SM6∼TM6B* flies were crossed with *UAS-CD8GFP, hs-flp; FRT40A, tub-Gal80; tub-Gal4/TM6B* flies.
<u>*tll*- MARCM clones:</u> *w;; FRT82B, tll[I49]/TM3,GFP,Ser* flies were crossed with *yw, hs-flp, UAS-GFP;; tub-Gal4, FRT82B, tub-Gal80/TM6C* flies.

### Antibody generation

Polyclonal antibodies were generated by Genscript (https://www.genscript.com/). The epitopes used for each immunization are listed below.

#### Erm

KTFSCLECGKVFNAHYNLTRHMPVHTGARPFVCKVCGKGFRQASTLCRHKIIHTSEKPHKCQTCGKAFNRSSTLNTHSRIHAGYKPFVCEYCGKGFHQKGNYKNHKLTHSGEKAYKCNICNKAFHQVYNLTFHMHTHNDKKPYTCRVCAKGFCRNFDLKKHMRKLHEIGGDLDDLDMPPTYDRRREYTRREPLASGYGQASGQLTPDSSSGSMSPPINVTTPPLSSGETSNPAWPRSAVSQYPPGGFHHQLGVAPPHDYPSGSAFLQLQPQQPHPQSQQHHQQQQRLSETFIAKVF

#### Ey

MFTLQPTPTAIGTVVPPWSAGTLIERLPSLEDMAHKDNVIAMRNLPCLGTAGGSGLGGIAGKPSPTMEAVEASTASHPHSTSSYFATTYYHLTDDECHSGVNQLGGVFVGGRPLPDSTRQKIVELAHSGARPCDISRILQVSNGCVSKILGRYYETGSIRPRAIGGSKPRVATAEVVSKISQYKRECPSIFAWEIRDRLLQENVCTNDNIPSVSSINRVLRNLAAQKEQQSTGSGSSSTSAGNSISAKVSVSIGGNVSNVASGSRGTLSSSTDLMQTATPLNSSESGGASNSGEGSEQEAIYEKLRLLNTQHAAGPGPLEPARAAPLVGQSPNHLGTRSSHPQLVHGNHQA LQQHQQQSWPPRHYSGSWYPTSLSEIPISSAPNIASVTAYASGPSLAHSLSPPNDIESLASIGHQRNCPVATEDIHLKKELDGHQSDETGSGEGENSNGGASNIGNTEDDQARLILK RKLQRNRTSFTNDQIDSLEKEFERTHYPDVFA

#### Esg

MHTVEDMLVEKNYSKCPLKKRPVNYQFEAPQNHSNTPNEPQDLCVKKMEILEENPSEELINVSDCCEDEGVDVDHTDDEHIEEEDEDVDVDVDSDPNQTQAAALAAAAAVAAAAAASVVVPTPTYPKYPWNNFHMSPYTAEFYRTINQQGHQILPLRGDLIAPSSPSDSLGSLSPPPHHYLHGRASSVSPPMRSEIIHRPIGVRQHRFLPYPQMPGYPSLGGYTHTHHHH

#### Hbn

MMTTTTSQHHQHHPIMPPAMRPAPVQESPVSRPRAVYSIDQILGNQHQIKRSDTPSEVLITHPHHGHPHHIHHLHSSNSNGSNHLSHQQQQQHSQQQHHSQQQQQQQQLQVQAKREDSPTNTDGGLDVDNDDELSSSLNNGHDLSDMERPRKVRRSRTTFTTFQLHQLERAFEKTQYPDVFTREDLAMRLDLSEARVQVWFQNRRAKWRKREKFMNQDKAGYLLPEQGLPEFPLGIPLPPHGLPGHPGSMQSEFWPPHFALHQHFNPAAAAAAGLLPQHLMAPHYKLPNFHTLLSQYMGLSNLNGIFGAGAAAAAAAASAGYPQNLSLHAGLSAMSQVSPPCSNSSPRESPKLVPHPTPPHATPPAGGNGGGGLLTGGLISTAAQSPNSAAGASSNASTPV SVVTKGED

#### Scro

MSSHGLAYTTRIERKSYRELQINRDQYFVTAPNEEDLVMSLSPKDTLIHTAISQHHQVDTSTKLNTNETSTQNTVSTAAAAAVAHHHHNLSSIHHLQNLHSQHQSTLFNSNH

#### Slp2

MVKIEEGLPSSEISAHSLHFQHHHHPLPPTTHHSALQSPHPVGLNLTNLMKMARTPHLK SSFSINSILPETVEHHDEDEEEDVEKKSPAKFPPNHNNNNLNTTNWGSPEDHEAESDPESDLDVTSMSPAPVANPNESDPDEVDEEFVEEDIECDGETTDGDAENKSNDGKPVKD KKGNE

#### Tll

MQSSEGSPDMMDQKYNSVRLSPAASSRILYHVPCKVCRDHSSGKHYGIYACDGCAGFFKRSIRRSRQYVCKSQKQGLCVVDKTHRNQCRACRLRKCFEVGMNKDAVQHERGPRNSTLRRHMAMYKDAMMGAGEMPQIPAEILMNTAALTGFPGVPMPMPGLPQRAGHHPAHMAAFQPPPSAAAVLDLSVPRVPHHPVHQGHHGFFSPTAAYMNALATRALPPTPPLMAAEHIKETAAEHLFKNVNWIKSVRAFTELPMPDQLLLLEESWKEFFILAMAQYLMPMNFAQLLFVYESENANREIMGMVTREVHAFQEVLNQLCHLNIDSTEYECLRAISLFRKSPPSASSTEDLANSSILTGSGSPNSSASAESRGLLESGKVAAMHNDARSALHNYIQRTHPSQPMRFQTLLGVVQLMHKVSSFTIEELFFRKTIGDITIVRLISDMYSQRKI

#### Otd

MAAGFLKSGDLGPHPHSYGGPHPHHSVPHGPLPPGMPMPSLGPFGLPHGLEAVGFSQGMWGVNTRKQRRERTTFTRAQLDVLEALFGKTRYPDIFMREEVALKINLPESRVQVWFKNRRAKCRQQLQQQQQSNSLSSSKNASGGGSGNSCSSSSANSRSNSNNNGSSSNNNTQSSGGNNSNKSSQKQGNSQSSQQGGGSSGGNNSNNNSAAAAASAAAAVAAAQSIKTHHSSFLSAAAAAASGGTNQSANNNSNNNNQGNSTPNSSSSGGGGGSQAGGHLSAAAAAAALNVTAAHQNSSPLLPTPATSVSPVSIVCKKEHLSGGYGSSVGGGGGGGGASSGGLNLGVGVGVGVGVGVGVSQDLLRSPYDQLKDAGGDIGAGVHHHHSIYGSAAGSNPRLLQPGGNITPMDSSSSITTPSPPITPMSPQSAAAAAHAAQSAQSAHHSAAHSAAYMSNHDSYNFWHNQYQQYPNNYAQAPSYYSQMEYFSNQNQVNYNMGHSGYTASNFGLSPSPSFTGTVSAQAFSQNSLDYMSPQDKYANMV

### Immunohistochemistry

Wandering third instar larval *Drosophila* optic lobes were fixed in 4% formaldehyde for 15 minutes at room temperature (with the exception of one staining using the mouse anti-eyeless, for which fixation was on ice for 30 minutes). After washing, they were incubated for 2 days with primary antibodies at 4°C. After washing the primary antibody, the brains were incubated with the secondary antibodies overnight at 4°C. The secondary antibodies were washed and the brains were mounted in Slowfade and imaged at a confocal microscope (Leica SP8) using a 63x glycerol objective. Images were processed in Fiji and Illustrator. At least four brains were analyzed for each experiment.

### Single-cell RNA seq

#### *Drosophila* optic lobes sample preparation

Developing central nervous systems from male and female flies were dissected from Canton-S wandering third instar larvae in PBS. The optic lobes were separated from the central brain using Vannas Spring Scissors with a 2mm cutting edge (Fine Science Tools Cat no. 15000-04). The optic lobes were dissociated into single cell suspension by incubating in 2mg/mL collagenase and 2mg/mL dispase in PBS for 15 minutes at 25C. The enzymes were then carefully removed and replaced with PBS + 0.1% BSA. The brains are soft but remain intact if pipetted slowly. The brains were pipetted up and down many times (> 100) until most large chunks of tissue are dissociated. The cells/tissue were kept cold by putting the tubes in ice. The cells were then filtered using 20um cell strainers. The concentration of the cell suspension was then measured staining the cells with 1/2000 Hoeschst, using an epifluorescent microscope and a 0.02-mm deep cytometer.

#### Library preparation and sequencing

Droplet-based purification, amplification and barcoding of single-cell transcriptomes were performed using Chromium Single Cell 3’ Reagent Kit v2 (10x Genomics) as described in the manufacturer’ s manual (Chromium Single Cell 3’ Reagent Kits v2 User Guide – Rev D), with a target recovery of 7,000 cells per experiment. We prepared 10 libraries, which were subjected to paired-end sequencing (26 × 8 × 98) with NovaSeq 6000 (Genome Technology Center at NYU Langone Health) to an average 50,000 reads per cell sequenced (that is, 350,000,000 reads for an experiment with 7,000 cells).

### Single-nucleus RNA seq

#### Human cortical plate sample preparation

Tissue was collected from de-identified prenatal autopsy specimens without neuropathological abnormalities under approved IRB protocol. The cortical plate was dissected fresh from the anterior frontal lobe of anatomically intact brain specimens with postmortem time interval less than 24 hours, and immediate fresh-frozen on dry ice.

#### Isolation and fluorescence-activated nuclear sorting (FANS) with hashing

All buffers were supplemented with RNAse inhibitors (Takara). 25mg of frozen postmortem human brain tissue was homogenized in cold lysis buffer (0.32M Sucrose, 5 mM CaCl_2_, 3 mM Magnesium acetate, 0.1 mM, EDTA, 10mM Tris-HCl, pH8, 1 mM DTT, 0.1% Triton X-100) and filtered through a 40 µm cell strainer. The flow-through was underlaid with sucrose solution (1.8 M Sucrose, 3 mM Magnesium acetate, 1 mM DTT, 10 mM Tris-HCl, pH8) and centrifuged at 107,000 g for 1 hour at 4°C. Pellets were re-suspended in PBS supplemented with 0.5% bovine serum albumin (BSA).

4 samples were processed in parallel. 2 million nuclei from each sample were pelleted at 500 g for 5 minutes at 4°C. Following centrifugation, nuclei were re-suspended in 100 µl staining buffer (2% BSA, 0.02% Tween-20 in PBS) and incubated with 1 µg of a unique TotalSeq-A nuclear hashing antibody (Biolegend) for 30 min at 4°C. Prior to FANS, volumes were brought up to 250 µl with PBS and DAPI (Thermoscientific) added to a final concentration of 1 µg/ml. DAPI positive nuclei were sorted into tubes pre-coated with 5% BSA using a FACSAria flow cytometer (BD Biosciences).

#### snRNAseq and library preparation

Following FANS, nuclei were subjected to 2 washes in 200 µl staining buffer, after which they were re-suspended in 15 µl PBS and quantified (Countess II, Life Technologies). Concentrations were normalized and equal amounts of differentially hash-tagged nuclei were pooled. A total of 40,000 (10,000 each) pooled nuclei were processed using 10x Genomics single cell 3’ v3 reagents. At the cDNA amplification step (step 2.2), 1 µl 2 µm HTO cDNA PCR “additive” primer was added^63^. After cDNA amplification, supernatant from 0.6x SPRI selection was retained for HTO library generation. cDNA library was prepared according to 10x Genomics protocol. HTO libraries were prepared as previously described^63^. cDNA and HTO libraries were sequenced at NYGC using the Novaseq platform (Illumina).

### Bioinformatic analyses

Detailed scripts and related R objects can be found here: https://drive.google.com/file/d/1geoY8AmtqFnJNi2OF7TlaJin8V9Nf-2C/view?usp=sharing

#### Mapping and integration of larval (L3) and pupal (P15) datasets

We mapped the sequenced libraries to the D. melanogaster genome assembly BDGP6.88 using CellRanger 3.0.1. We kept only genes that were expressed in at least 3 cells across all cells and cells with counts for at least 200 genes for further analysis. After processing, the dataset comprised 49,893 cells passing quality filters, with a median of 3,635 UMIs and 1,343 genes per cell.

We used the procedure implemented in Seurat v.3 to remove batch effects from our sequenced libraries. We used default parameters except for the dimensionality for which we tried the values 100, 150 and 200. We compared the results using the Seurat function LocalStruct with default parameters. The results obtained were 83.7%, 84.9% and 83.6%, respectively. We therefore chose a dimensionality of 150 for the larval dataset.

The dataset was then clustered with a resolution of 2. Notably, in this developing structure, cells are clustered both by identity and by differentiation stage. For example, Mi1 cells fall into 2 clusters, an immature (cluster 23) and a mature cluster (cluster 53).

Larval and pupal datasets were merged using default parameters.150 PCs were used subsequently for generating the UMAP to remain consistent with the integration of the larval dataset.

#### Spatial patterning analysis

To focus on the heterogeneity within the neuroepithelial cells, the larval dataset is further subsetted using marker expression with Seurat v3. Expression of neuroepithelial markers *shg*, *tom*, and *brd* were examined for each cluster^64^. Clusters with average expression higher than 95^th^ percentile of normalized expression of Tom and Brd were selected as neuroepithelial clusters. DE-Cadherin (Shg) is known to be enriched in neuroepithelial cells^65^ and is enriched in the selected clusters (logFC = 0.75, adjusted p value = 0).

Principal components were calculated using variable features found in the subsetted neuroepithelial cells. Examination of PC1 revealed Tll, an early marker of lamina precursor cells^66^, is expressed in a near-mutually exclusive fashion with Hth (enriched in neuroepithelium and young medulla neuroblasts), suggesting the subset contained both OPC neuroepithelium and lamina precursor cells. To keep only OPC neuroepithelial, we sub-clustered the cells and examined the average expression of Hth and Tll (enriched in lamina precursor cells) for each cluster. This process is performed iteratively to keep only Hth+/Tll-clusters. The remaining cells were assigned as OPC neuroepithelium for further analysis of spatial temporal factors.

#### Trajectory analysis: identification of candidate tTFs

To study temporal patterning in neuroblasts, we first identified the cluster that corresponded to the medulla neuroblasts (cluster 9) based on the expression of Dpn, as well as the expression of the known temporal factors. We extracted the counts from these cells and inputted them into Monocle. We used default parameters to order the cells in pseudotime. We used the DDRTree method for dimensionality reduction. The cells were then ordered in pseudotime and the beginning and end of the trajectory were defined based on the expression of the known tTFs (i.e. Hth marked the beginning of the trajectory and Tll marked the end). We then looked at the expression along the pseudotime of 629 genes annotated as transcription factors in FlyBase to identify the candidate tTFs.

#### Merging of larval and pupal Mi1 and DE analysis over pseudotime

Larval and pupal (P15, P30, P40, P50, and P70) datasets were merged after cells were batch corrected for each stage separately. Standard Monocle workflow was followed to generate trajectories. The L3 and P15 trajectories were ordered manually.

Based on the way the developing optic lobe develops, there are cells at the same differentiation stage in the L3 and P15 datasets. We, therefore, decided to align these two datasets in order to get a continuum of expression. We tested different genes and ended up using “Ggamma30A” as a reference gene. Ggamma30A starts increasing in the middle of the L3 trajectory and continues all the way to P15 in linear manner. We adjusted the expression of Ggamma30A in P15 using linear regression, which was then applied to all genes of P15. This does not change the dynamics of expression, just the relative levels, and serves the purpose of aligning the trajectories over pseudotime of L3 and P15.

To identity differentially expressed genes along the differentiation trajectory from L3 to P70, we used two methods: “principal graph” and “knn”. We selected genes that were identified as differentially expressed with at least one of the two methods. We then used the find_gene_modules function to group the differentially expressed genes into modules of genes that co-vary. These genes were then used for GO analysis.

#### GO enrichment analysis

We performed GO enrichment analysis and calculated enrichment for ‘Biological Process’ using The Gene Ontology Resource (http://geneontology.org/) using a Fisher’s exact test to calculate p-value. Multiple testing correction was performed by calculating the False Discovery Rate.

To find the expression of GO terms over time, we added and normalized the expression of all genes that belong to a specific GO term and plotted it over pseudotime or on the UMAP.

#### Analysis of human data

We mapped the sequenced libraries to the H. sapiens genome assembly GRCh38 (hg38) using CellRanger 3.1.0. For the hashtag oligos (HTO), we used the CITE-seq-Count 1.4.2 version to align HTO to 10x barcodes using the following command:

CITE-seq-Count -R1 reas1 -R2 read2 -T 1 -t tag -cbf 1 -cbl 16 -umif 17 -umil 26 -cells 40000 -o output --sliding-window # --dense

After processing, the dataset comprised 3,363 cells passing quality filters, with a median of 4,736 UMIs and 2,414 genes per cell.

We selected the radial glia (expressing Pax6), intermediate progenitors (expressing Eomes), and neurons (expressing NeuroD2) that were forming a trajectory in UMAP and imported the data into Monocle and used default parameters to calculate the trajectories. We used the find_gene_modules function to group genes into 6 modules of genes that co-vary. These modules were then used for GO analysis.

#### Analysis of mouse cortical data

The dataset that was generated by Telley et al.^18^ was downloaded from GEO (GSE118953). The raw counts were inputted into Seurat and the standard workflow was followed (log-normalization, followed by clustering and UMAP using 25 PCs, and clustering was done with a resolution of 2). The radial glia clusters (clusters 2 and 3) were identified based on the expression of known radial glia markers, such as SOX2 and PAX6. Radial glia from different embryonic days 12, 13, 14, and 15 were used to generate the violin plots of Extended Data Figure 7A.

## Acknowledgements

We are indebted to the fly community for gifts of antibodies and fly lines; Cheng-Yu Lee, Deborah Hursh, John Nambu, Andrew Tomlinson, and Mitshuhiko Kurusu for fly lines, Peter Gergen, Kwangwook Choi, Hermann Aberle, and Dorothea Godt for antibodies. We thank Isabel Holguera, Sergio Cordoba, Chris Doe, Stein Aerts, and Tzumin Lee for constructive feedback on the manuscript. We thank Sergio Cordoba for the optic lobe illustrations and Kazi Hossain and Albert Tadros for help with preliminary experiments. Finally, we thank all members of the Desplan Lab for helpful discussions. This work was supported by NIH grant EY019716 to C.D. N.K. was supported by the National Eye Institute (K99 EY029356-01). A.M.R was partly supported by funding from NIH (T32 HD007520), and by NYU’s GSAS MacCracken Program and a Dean’s Dissertation Fellowship. A.M.J. was supported by the NYU SURP program. M.N.O is a Leon Levy Neuroscience Fellow. F.S. and Y.C.C. are supported by the New York University (MacCracken Fellowship).

## Author contributions

N.K., A.M.R., and C.D. designed the project. N.K. A.M.R., A.E., and T.T performed the genetic experiments. N.K., A.M.R., A.E., L.D., T.T, A.M.J., and Y.C.C., analyzed the data. N.K., M.N.O., and F.S. acquired the fly scRNA-seq data. N.M.T., J.F.F., Z.S, and P.R. acquired the human scRNA-seq data. U.W. generated the *hbn* mutant. N.K., L.D., A.M.J., and Y.C.C. performed scRNA-seq data analysis. N.K., A.M.R., and C.D. wrote the manuscript. All authors edited the manuscript.

## Declaration of Interests

Authors declare no conflicts of interest.

## Data Availability

All *Drosophila* raw and processed data referenced were uploaded to GEO: accession number GSE167266.

The human source data described in this manuscript are available via the PsychENCODE Knowledge Portal (https://psychencode.synapse.org/). The PsychENCODE Knowledge Portal is a platform for accessing data, analyses, and tools generated through grants funded by the National Institute of Mental Health (NIMH) PsychENCODE program. Data is available for general research use according to the following requirements for data access and data attribution: (https://psychencode.synapse.org/DataAccess).

## Extended Data figure legends

**Extended Data figure 1:**
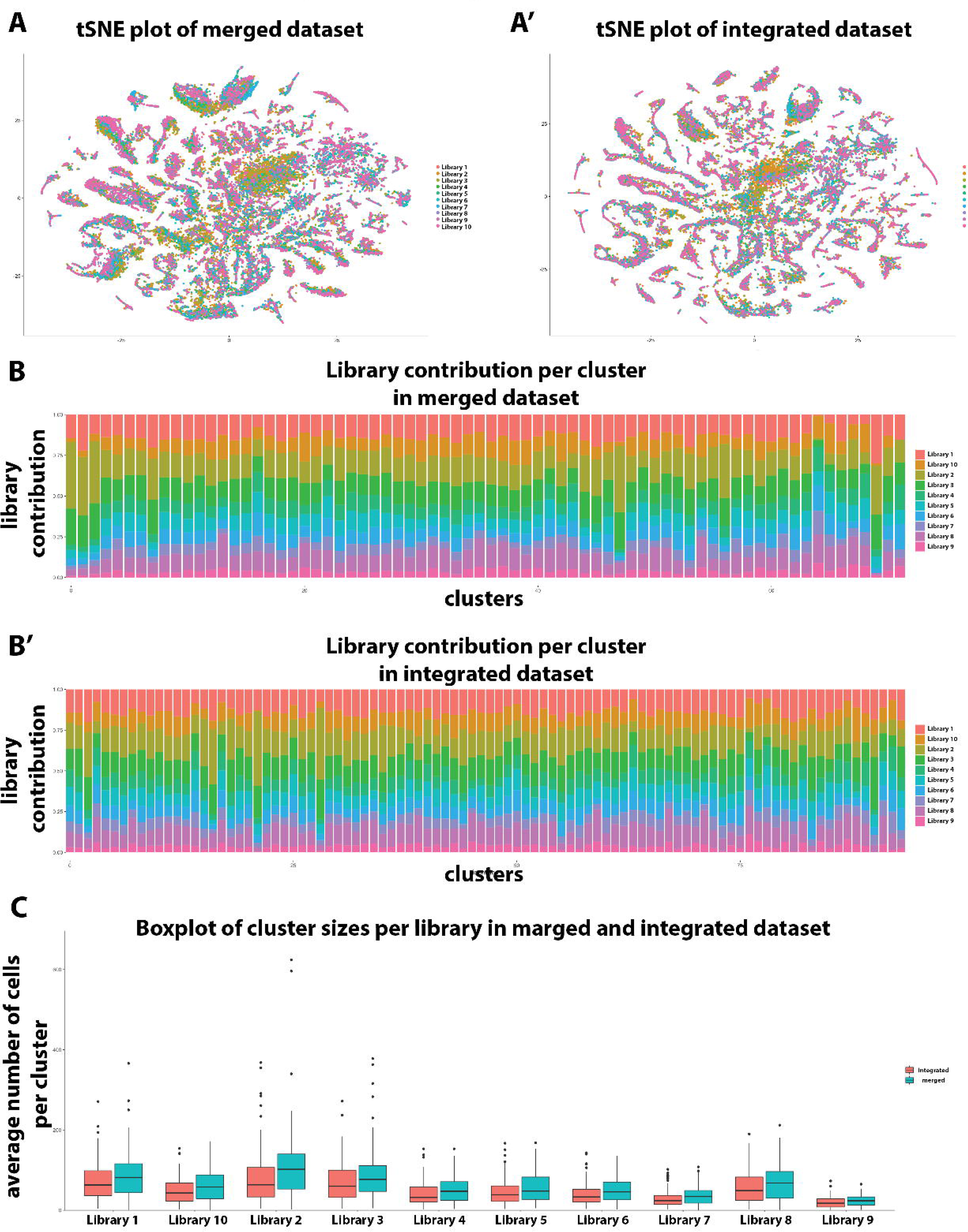
Integration of libraries. (A-A’) Comparison of library distribution on a tSNE plot of datasets before (merged) and after library integration (batch effect correction). Before integration, there is a clear bias in the distribution of the libraries within clusters. After integration, this bias is largely eliminated. (B-B’) Comparison of the library contribution in each cluster before (merged) and after library integration (batch effect correction). With the exception of few clusters in each case, all clusters have a similar percentage of cells coming from each library. (C) Comparison of cluster sizes per library before (merged) and after library integration (batch effect correction). The variance in the merged dataset is larger than the one in the integrated one, indicating noise that was potentially alleviated by the batch correction.

**Extended Data figure 2:**
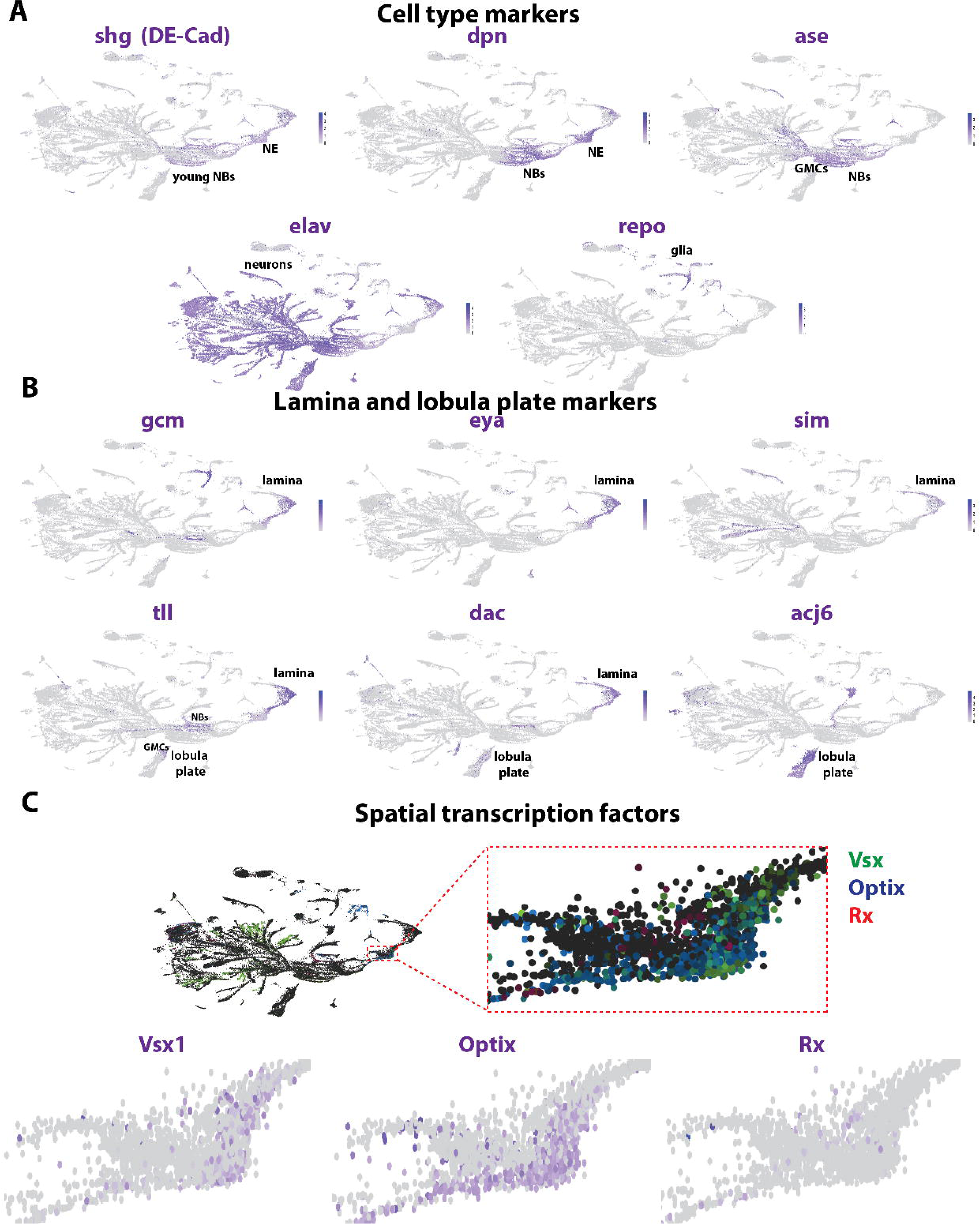
Annotation of single-cell sequencing UMAP plot. (A) UMAP plots showing the expression of different cell type markers that allows for the annotation of the different clusters. *Shotgun/DE-Cad (shg)* is expressed in the neuroepithelium (NE) and young neuroblasts (NBs). *Deadpan (dpn)* is expressed in neuroepithelium and neuroblasts. *Asense (ase)* is expressed in the neuroblasts and GMCs. *Elav* is mostly expressed in the neurons, although the transcript can already be seen in the GMCs. Finally, *repo* is expressed in glial cells. (B) Expression of markers for the lamina and the lobula plate. Lamina is marked by the expression of *gcm, eya, sim, tll,* and *dac,* while lobula plate expressed strongly *acj6,* faintly *dac,* and the progenitors (NBs and GMCs) express *tll*. (C) (Left). Spatial transcription factors (*Vsx, Optix,* and *Rx*) are not expressed in medulla neuroblasts, while only Vsx is expressed in some neuronal types (it is unknown whether this expression reflects their origin from the Vsx spatial domain). (Right) sTFs are only expressed in the neuroepithelium in largely non-overlapping domains. (Bottom). UMAP plots with the expression of individual sTFs.

**Extended Data figure 3:**
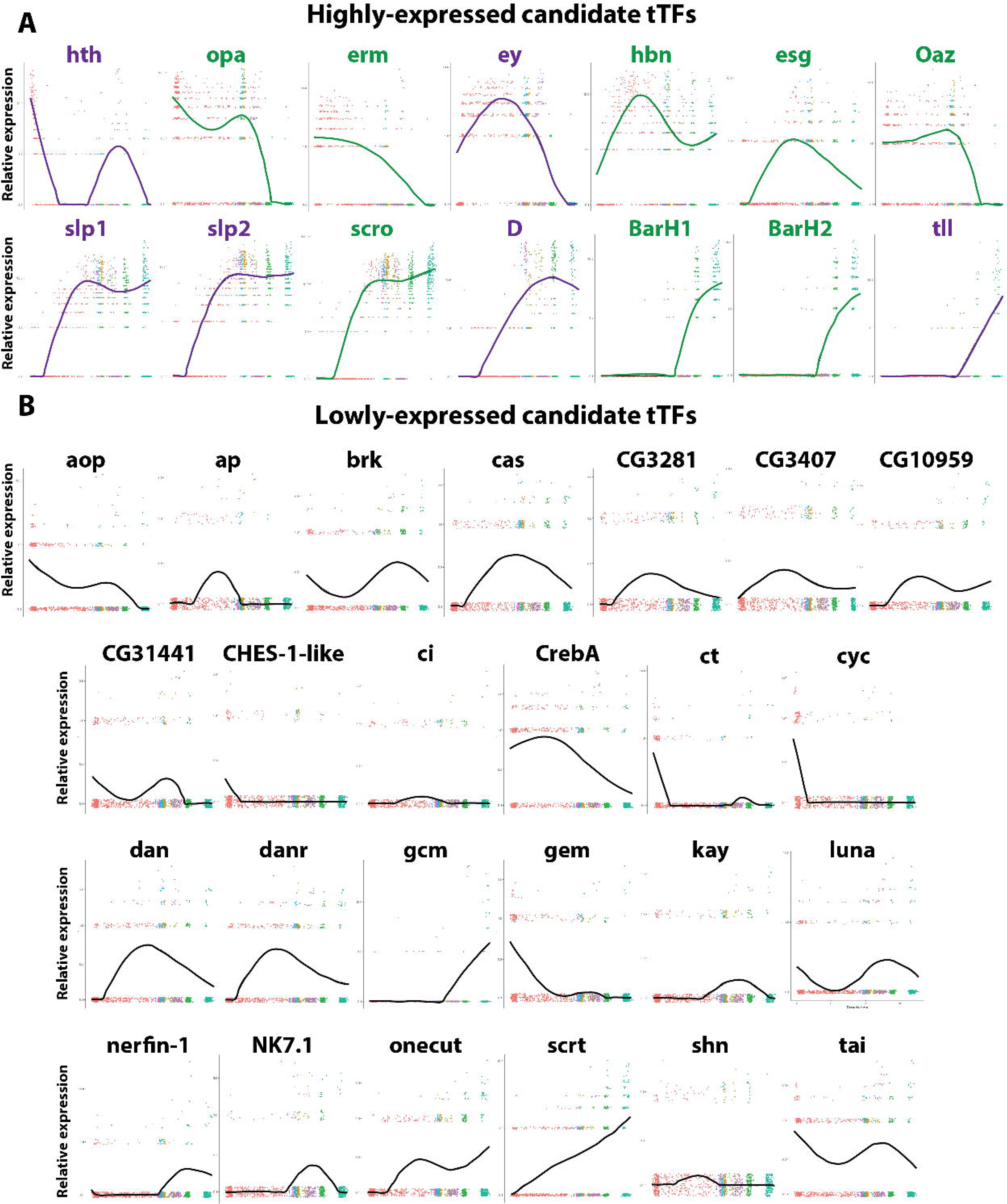
Candidate tTF expression in neuroblast trajectory. Expression pattern over pseudotime of all TFs that were found to be expressed in a temporal manner along the neuroblast trajectory. (A) 14 transcription factors were found to be expressed temporally in high relative expression levels. These include the already known tTFs (*hth*, *ey*, *slp1*, *slp2*, *D*, and *tll* in green), as well as eight new candidate tTFs (in purple). (B) Another 25 transcription factors were found to be expressed temporally in lower relative expression levels. *Ap, ct, gcm*, and *gem* expression was tested in developing optic lobes and they were not expressed temporally, so these 25 transcription factors were excluded from downstream analysis.

**Extended Data figure 4:**
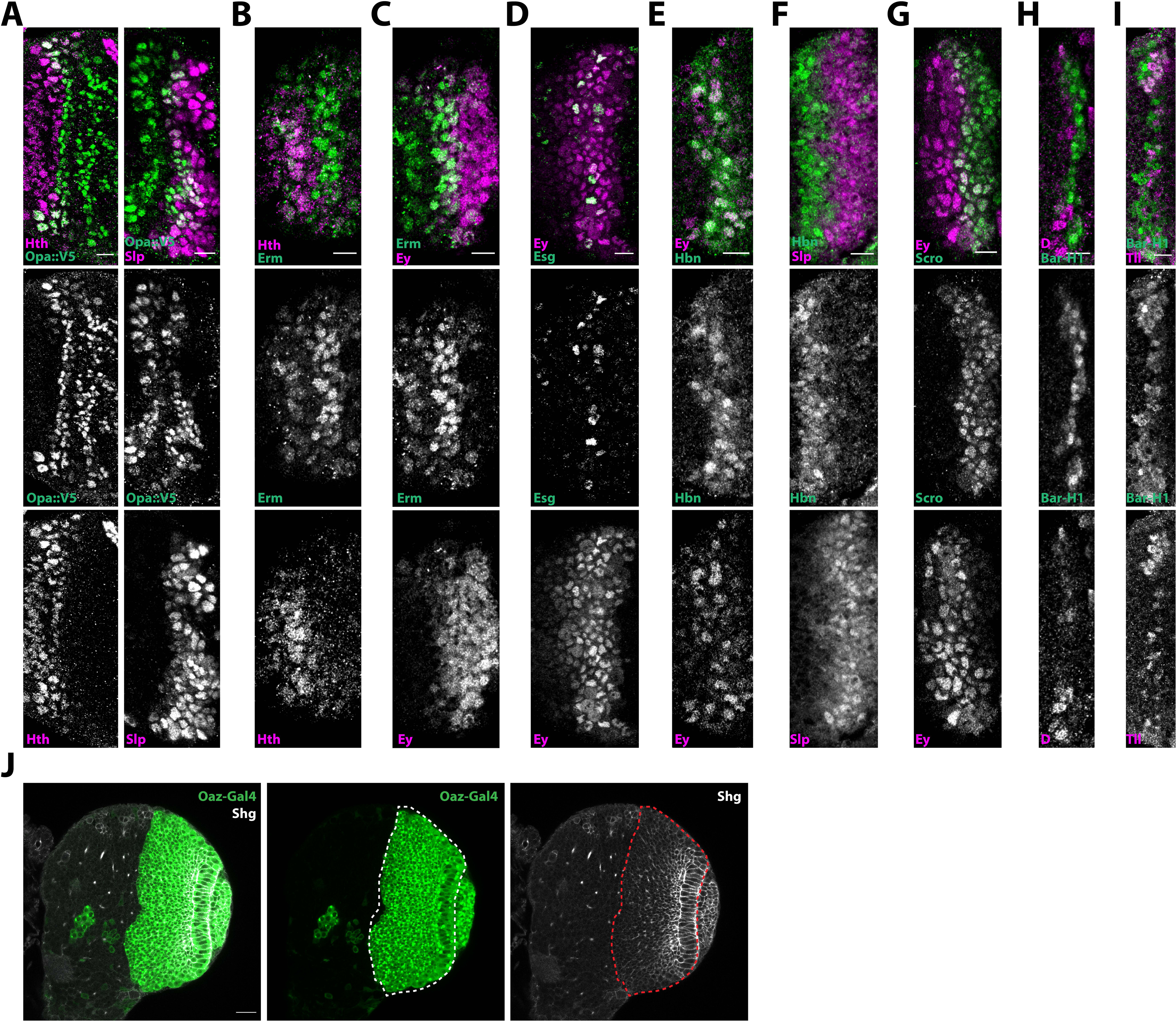
Expression of candidate tTFs in relation to the known tTFs. (A-I) Antibody stainings (including single channel images) of newly identified temporal transcription factors (green) and previously known ones (purple) show that the candidate temporal transcription factors are indeed expressed temporally. Scale bar: 10um (A) Opa is expressed in two waves, one succeeding and partially overlapping (arrow) with the Hth window and one immediately before Slp. (B) Erm is expressed immediately after Hth. (C) Erm starts being expressed before Ey, partially overlapping with it (D) Esg is expressed in a salt-and-pepper manner within the Ey temporal window. (E) Hbn expression is nested within Ey temporal window. (F) Hbn is expressed before Slp1. (G) Scro is expressed immediately after Ey. (H) BarH1 is expressed after the D temporal window. (I) BarH1 is expressed right before Tll. (J) Oaz-Gal4 driving UAS>GFP expression (green). The transgene is expressed in the neuroepithelium and all neuroblasts of the developing medulla (dashed line). The neuroepithelium and neuroblasts are marked by Shg (white). Scale bar: 20um

**Extended Data figure 5:**
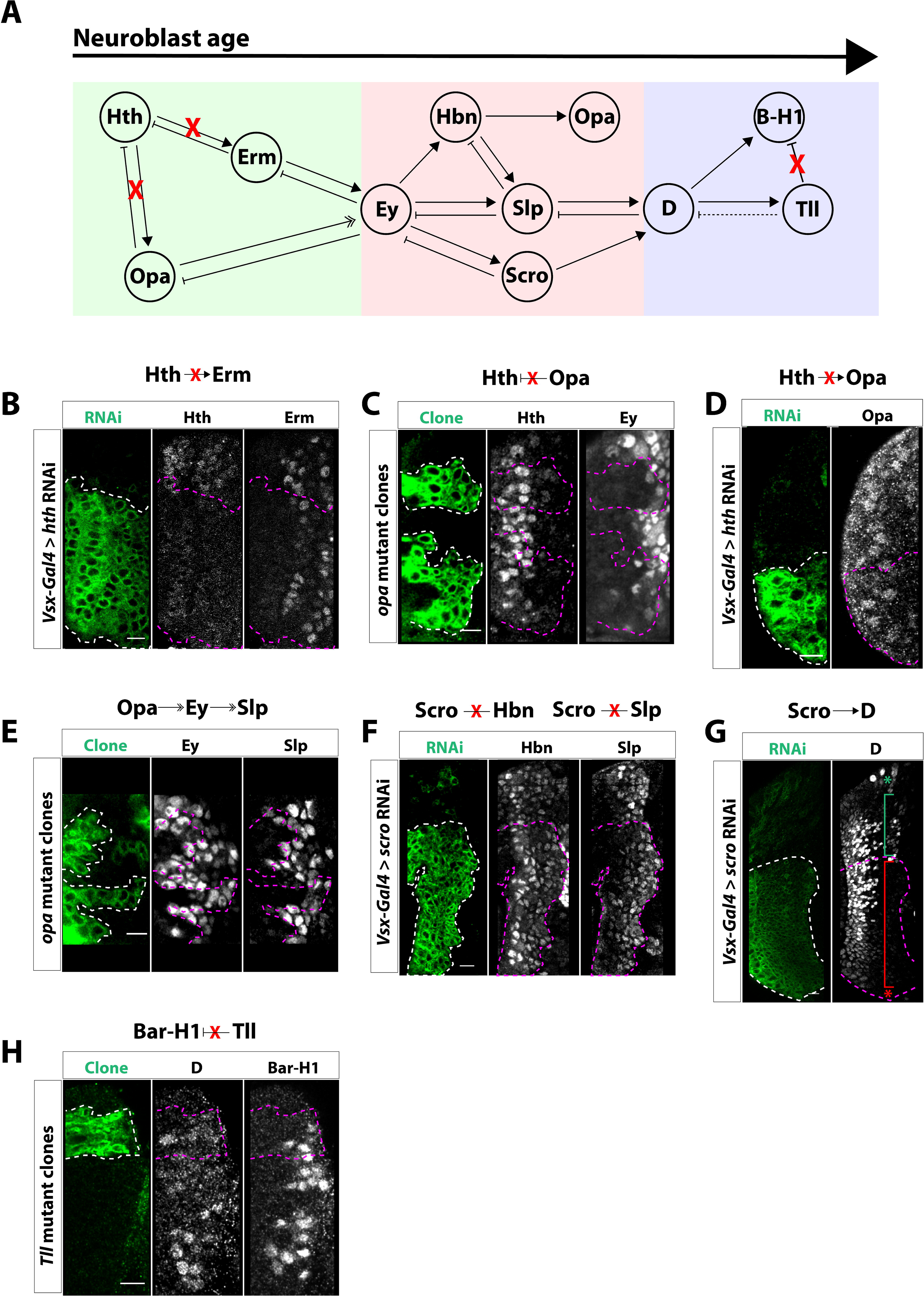
Negative genetic interaction between tTFs. (A) Diagram of genetic interactions between tTFs in medulla neuroblasts. Red “X”’s within the diagram indicate no genetic interaction. Within the early unit (green box), we identified two new tTFs: Odd paired (Opa) and Earmuff (Erm). Homothorax (Hth) does not activate Erm or Opa. Furthermore, Opa does not repress Hth. Within the middle unit (red box), we identified three new temporal factors: Homeobrain (Hbn), Scarecrow (Scro) and Opa. No genetic interactions exist between Hbn and Scro or between Sloppy paired (Slp) and Scro. Within the late unit (blue box) we identified one new temporal factor: BarH1 (B-H1). Tailless (Tll) does not inhibit BarH1. (B) In cells expressing *hth* RNAi driven by *Vsx-Gal4* (GFP: green), Erm expression is not affected, indicating that Hth does not activate Erm. (C) In *opa* mutant clones (GFP: green), Hth is not affected and Eyeless (Ey) expression is delayed, demonstrating that Opa does not inhibit Hth and only helps to time the expression of Ey. (D) Furthermore, no positive genetic interaction exists between Hth and Opa, as cells expressing *hth* RNAi (GFP: green) maintain Opa expression. (E) As expected, in *opa* mutant clones (GFP: green), not only is Ey expression delayed but Slp is also delayed. (F) Loss of Scro following the expression of *scro* RNAi (GFP:green) does not affect the expression of Homeobrain (Hbn) or Slp, as might be expected given their normal coexpression (see Figure 2). (G) To support our observation that Scro activates D, knocking down *scro* by expressing *scro* RNAi (GFP:green) leads to the loss of D expression in neurons born during the D temporal window (red bracket). The green bracket indicates D+ neurons born from neuroblasts not expressing *scro* RNAi. The red and green asterisks indicate neuroblasts at the tips of epithelium. In the GFP+ region (RNAi expressing neuroblasts) D is lost. The D+ neurons to the left of the image are those born from the Ey temporal window. (H) In *Tll* mutant clones (GFP:green), neither D or BarH1 expression are affected, indicating that Tll is not necessary to inhibit either factor. Scale bar: 10um

**Extended Data figure 6:**
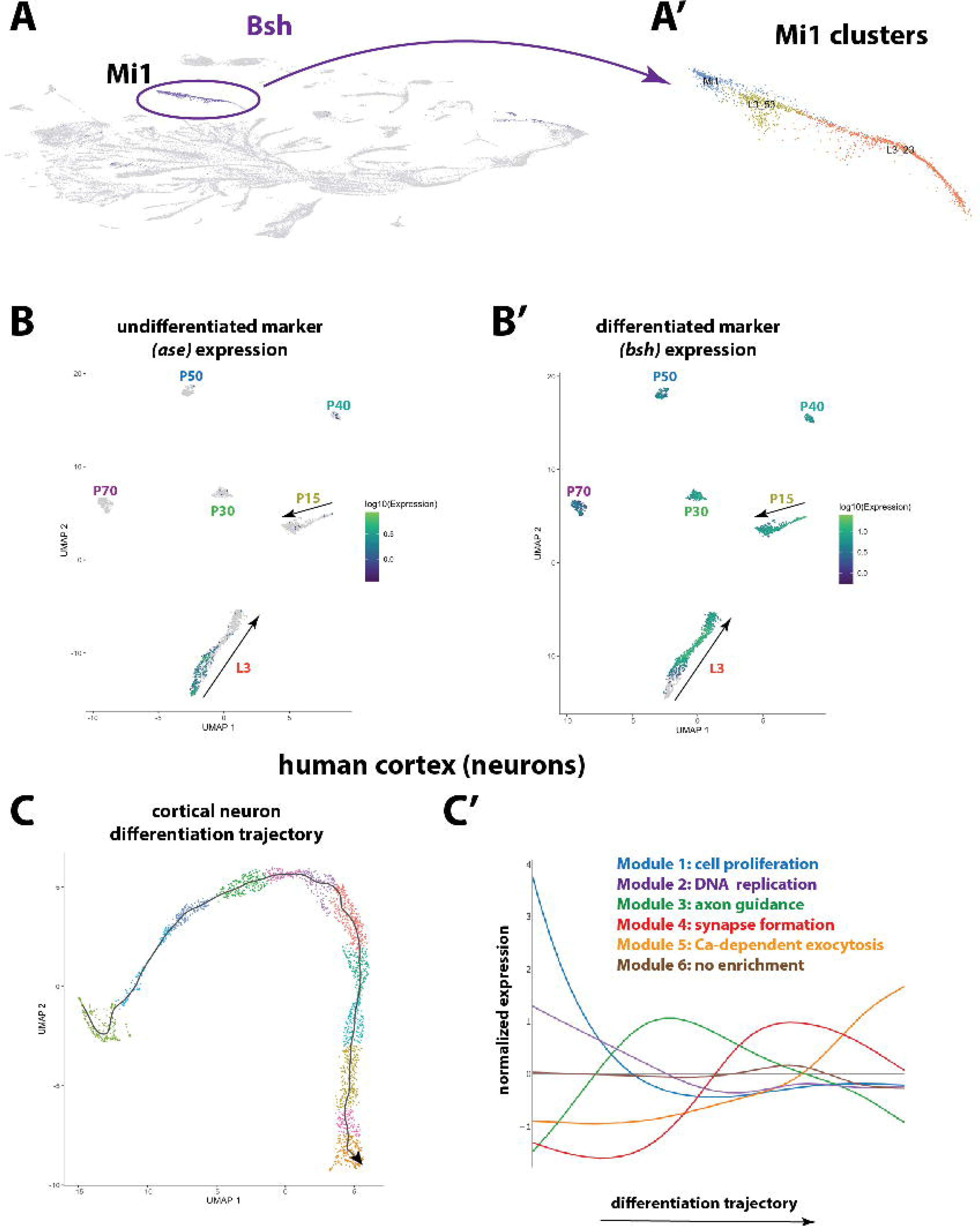
Neuronal differentiation in flies and humans. (A) *Bsh* is expressed almost exclusively in Mi1s and was used to identify the Mi1 clusters. (A’) Cluster Mi1 represents the pupal annotated cluster. Cluster L3_23 consists of GMCs that give rise to Mi1s and newly-born Mi1s, while cluster L3_53 is more mature Mi1 cells, as assessed by their proximity to the P15 Mi1 cells. (B) UMAP plot of Mi1 cells at different stages of differentiation from L3 to P70. The expression of *ase* (B) and *bsh* (B’) were used to find the beginning and end, respectively, of the L3 trajectory. (C) UMAP plot showing the trajectory of 3,363 single-cell transcriptomes of the developing human cortex (gestational week 19), as generated by Monocle3. The orientation of the trajectory was identified by looking at the expression of marker genes for progenitors, intermediate progenitors and neurons (see Figure 5F). (C’) Differential expression analysis along the trajectory of the cortical neurons identified six modules of genes. Gene Ontology enrichment analysis found the first two modules to be enriched in terms, such as cell proliferation and DNA replication; they likely correspond to the progenitor cells. Then, the third module is enriched in neurite development terms, such as axon guidance, while the fourth one is enriched in terms related with synapse formation. The fifth one contains “functional genes”, such as calcium-dependent exocytosis. The sixth module does not show a clear peak of expression and no GO terms were found to be enriched.

**Extended Data Figure 7:**
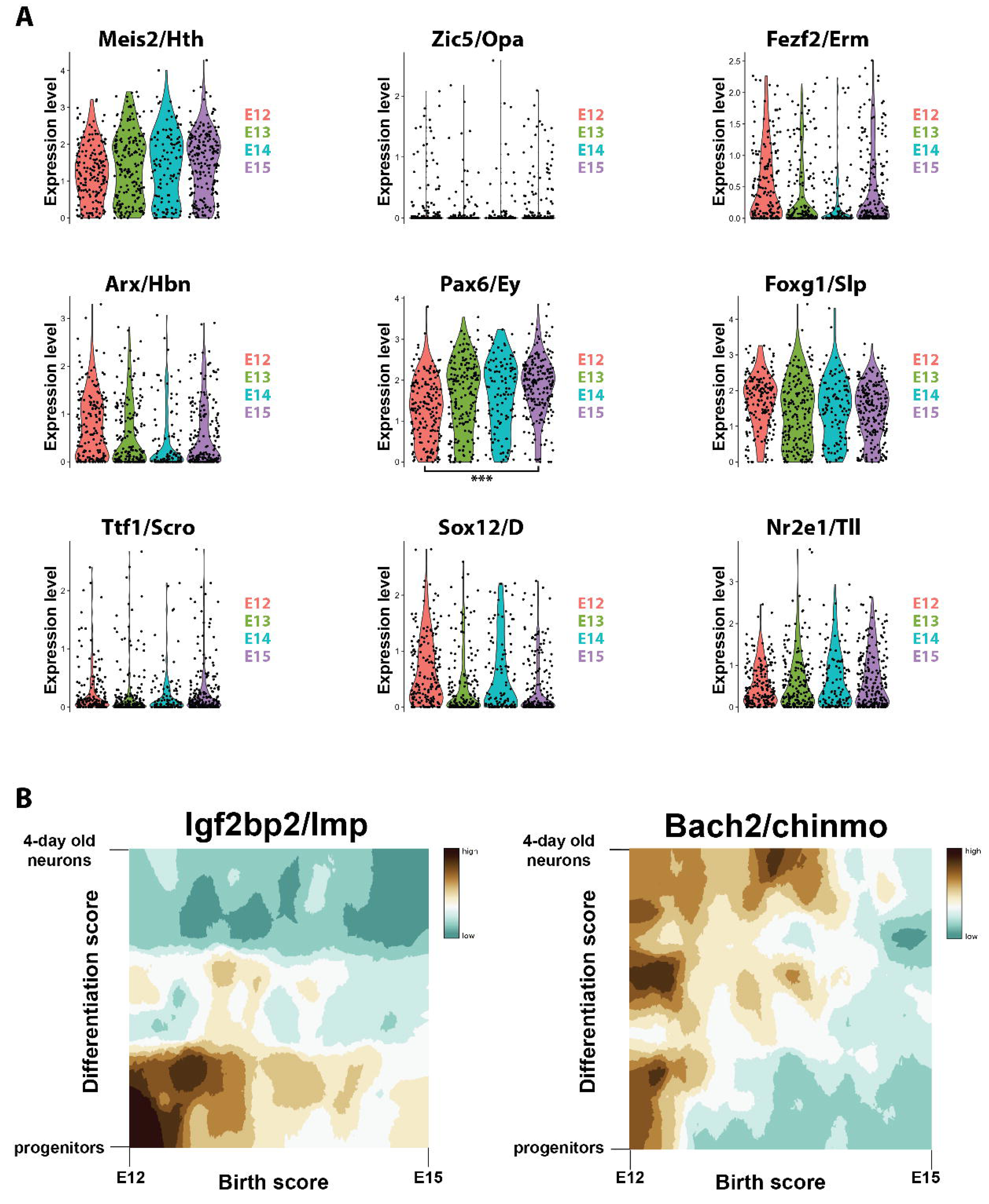
Expression of Drosophila tTFs in mouse cortical radial glia. (A) The mouse orthologs of the *Drosophila* optic lobe temporal transcription factors are not expressed in temporal windows in mouse cortical radial glia during embryonic stages E12-E15, which span their neurogenic period, with the exception of Pax6, which is enriched in young progenitors (adjusted p-value = 1.969240e-08). (B) Heatmap of expression of Igf2bp1 and Bach2 (orthologs of *Drosophila* Imp and Chinmo, respectively) in radial glia and neuronal progeny. Igf2bp1 is expressed in young apical progenitors, while Bach2 is expressed in young apical progenitors and neurons that are born by these young progenitors. Source: http://genebrowser.unige.ch/telagirdon/

**Supplementary Table 1:** Temporal and neurotransmitter identity of all medulla cell types. This table shows the predicted temporal identity of every medulla neuronal cell types (i.e. the temporal window at which these neurons were generated); predictions are made using the expression of tTFs or published tTF targets and the position of the cluster in the UMAP. Moreover, it shows the neurotransmitter identity of all clusters at L3 and adult stages.

**Supplementary Table 2:** Fly strains and antibodies.

## Notes

### Competing Interest Statement

The authors have declared no competing interest.

