## Supplemental Table 2 for "A comprehensive series of temporal transcription factors in the fly visual system"

| REAGENT or RESOURCE | SOURCE | IDENTIFIER |
| --- | --- | --- |
| Fly strains | |  |
| Drosophila, Canton S | Bloomington Drosophila Stock Center (BDSC) | 64349 |
| Drosophila, Opa:V5 | (Abdusselamoglu et al, 2019) | N/A |
| Drosophila, MzVum-Gal4 | BDSC | 29031 |
| Drosophila, Oaz-Gal4 | BDSC | 80596 |
| Drosophila, hth-RNAi | Vienna Drosophila Resource Center (VDRC) | v100630 |
| Drosophila, scro-RNAi | BDSC | 33890 |
| Drosophila, erm-FRT40A | (Weng et al, 2010)  Gift from Cheng-yu Lee | N/A |
| Drosophila, opa-FRT82B | (Lee et al, 207)  Gift from Deborah Hursh | N/A |
| Drosophila, ey-FRT80B | (Li et al, 2013)  Desplan lab | N/A |
| Drosophila, ey rescue-FRT80B | (Li et al, 2013)  Desplan lab | N/A |
| Drosophila, D-FRT2A | (Nambu et al, 1996)  Gift from John Nambu | N/A |
| Drosophila, hbn-FRT42B | (Kolb et al, 2021)  Walldorf lab | N/A |
| Drosophila, slp-FRT40A | (Sato et al, 2007)  Gift from Andrew Tomlinson | N/A |
| Drosophila, tll-FRT82B | (Pignoni et al, 1990)  Gift from Mitsuhiko Kurusu | N/A |
| Drosophila, MARCM FRT82B | BDSC | 86311 |
| Drosophila, FRT2A | (Li et al, 2013)  Desplan lab | N/A |
| Drosophila, MARCM FRT42B | Uwe Walldorf | N/A |
| Drosophila, MARCM FRT40A | (Li et al, 2013)  Desplan lab | N/A |
| Antibodies |  |  |
| Sheep anti-GFP (1:200) | BioRad | 4745-1051 |
| Chicken anti-GFP (1:1000) | Millipore Sigma | 06-896 |
| Chicken anti-V5 (1:500) | Abcam | ab9113 |
| Rat anti-elav (1:20) | Developmental Studies Hybridoma Bank (DSHB) | 7E8A10 |
| Rat anti-shg (1:100) | DSHB | DCAD2 |
| Guinea pig anti-Hth (1:500) | (Ozel, Simon et al, 2020)  Desplan lab | N/A |
| Rabbit anti-Opa (1:500) | (Mendoza-Garcia et al, 2017)  Gift from Peter Gergen | N/A |
| Rat anti-Erm (1:100) | This paper | N/A |
| Rabbit anti-Ey (1:250) | This paper | N/A |
| Mouse anti-Ey (1:10) | DSHB | Anti-eyeless |
| Rat anti-Esg (1:200) | This paper | N/A |
| Rabbit anti-Hbn (1:1000) | This paper | N/A |
| Guinea pig anti-Scro (1:100) | This paper | N/A |
| Guinea pig anti-Slp2 (1:200) | This paper | N/A |
| Rabbit anti-D (1:500) | modEncode | N/A |
| Rabbit anti-Bar-H1 (1:500) | Gift from Kwangwook Choi | N/A |
| Guinea pig anti-Tll (1:500) | This paper | N/A |
| Guinea pig anti-Tj (1:250) | (Gunawan et al, 2013)  Gift from Dorothea Godt | N/A |
| Guinea pig anti-Otd (1:1000) | This paper | N/A |
| Guinea pig anti-Cha (1:400) | (Konstantinides et al, 2018)  Desplan lab | N/A |
| Guinea pig anti-VGlut (1:1000) | (Mahr and Aberle, 2006)  Gift from Hermann Aberle | N/A |
| Donkey anti-guinea pig Alexa 405 (1:100) | Jackson ImmunoResearch | 706-475-148 |
| Donkey anti-chicken Alexa 405 (1:100) | Jackson ImmunoResearch | 703-475-155 |
| Donkey anti-rat Alexa 405 (1:100) | Jackson ImmunoResearch | 712-475-153 |
| Donkey anti-sheep Alexa 488 (1:500) | Jackson ImmunoResearch | 713-545-147 |
| Donkey anti-chicken Alexa 488 (1:500) | Jackson ImmunoResearch | 703-545-155 |
| Donkey anti-rat Alexa 488 (1:500) | Jackson ImmunoResearch | 712-545-153 |
| Donkey anti-mouse Alexa 488 (1:500) | Jackson ImmunoResearch | 715-545-151 |
| Donkey anti-rabbit Alexa 488 (1:500) | Jackson ImmunoResearch | 711-545-152 |
| Donkey anti-guinea pig Alexa 488 (1:500) | Jackson ImmunoResearch | 706-545-148 |
| Donkey anti-rabbit Cy3 (1:500) | Jackson ImmunoResearch | 711-165-152 |
| Donkey anti-mouse Cy3 (1:500) | Jackson ImmunoResearch | 715-165-150 |
| Donkey anti-rat Cy3 (1:500) | Jackson ImmunoResearch | 712-165-153 |
| Donkey anti-rat Alexa 647 (1:200) | Jackson ImmunoResearch | 712-605-153 |
| Donkey anti-guinea pig Alexa 647 (1:200) | Jackson ImmunoResearch | 706-605-148 |
| Donkey anti-rabbit Alexa 647 (1:200) | Jackson ImmunoResearch | 711-605-152 |
